# Improved Immune Responses and Tuberculosis Protection by Aerosol Vaccination with recombinant BCG expressing ESX-1 from *Mycobacterium marinum*

**DOI:** 10.1101/2025.09.18.677039

**Authors:** Fadel Sayes, Wafa Frigui, Alexandre Pawlik, Cécile Tillier, Magali Tichit, David Hardy, Roland Brosch

## Abstract

The currently licensed anti-tuberculosis (TB) vaccine, *Mycobacterium bovis* BCG, provides limited protection against pulmonary TB in adolescents and adults, the main cause of TB transmission and mortality. To obtain an improved BCG-based vaccine candidate with increased immune signaling but still low virulence, we have previously generated a recombinant BCG strain named BCG::ESX-1*^Mmar^*, which is heterologously expressing ESX-1 functions of *Mycobacterium marinum* and thereby modulates the host innate immune responses via phagosomal rupture-associated induction of type I interferon production and enhanced inflammasome activity, leading to superior protection against TB disease in murine infection models. As protection may also vary with the route of vaccination, here, we have explored aerosol vaccination relative to subcutaneous vaccination, using BCG Pasteur and BCG::ESX-1*^Mmar^*. We found that mice vaccinated via the aerosol route with BCG Pasteur or BCG::ESX-1*^Mmar^* both yielded higher frequencies of CD4^+^ and CD8^+^ T effector memory (TEM) cells in the lungs compared to subcutaneously immunised mice, whereas comparable poly-functional Th1 (IL-2, TNF-α and IFN-γ) cytokine-producing subsets were observed in the spleens of all vaccinated mice. Significantly higher IL-17 responses without severe lung pathology were seen in the lungs of aerosol-vaccinated mice associated to local and transient inflammatory cytokine responses and immune cell infiltrations. Aerosol vaccination also elicited high amounts of humoral IgG and IgM responses in the bronchoalveolar lavage fluid and induced substantial lung CD69^+^ CD103^+^ T resident memory (TRM) cells, containing both CD4^+^ and CD8^+^ T subsets, in the airways of immunised mice, whereas this was not the case for subcutaneous vaccination. These effects led to significant improved protection against *M. tuberculosis* and reduced lung pathology in aerosol-vaccinated mice compared to subcutaneously vaccinated mice. Moreover, BCG::ESX-1*^Mmar^* induced enhanced T-cell immunity and superior protection compared to parental BCG Pasteur for both vaccination routes and thereby represents an interesting candidate for developing improved vaccination strategies against TB.

**Author summary:** Anti-tuberculosis vaccine efficacy is influenced by multiple parameters, including the immunogenicity of the vaccine strain, the type of preclinical host model used, and the route of vaccination. Given recent advances in the field of mucosal vaccination, in the current study we were particularly interested to explore and compare aerosol-based vaccination with standard subcutaneous vaccination in a C57BL/6J mouse model using our recently developed recombinant BCG::ESX-1*^Mmar^* vaccine candidate in comparison with parental BCG Pasteur. Our results show that in this setting the protective efficacy of mucosal vaccination was superior to subcutaneous vaccination for both vaccine strains, whereby the use of BCG::ESX-1*^Mmar^* induced additional benefits in terms of bacterial load reduction compared to standard BCG Pasteur. Taken together, we propose that aerosol vaccination using BCG::ESX-1*^Mmar^* as live-attenuated vaccine candidate is a promising and powerful combination for obtaining improved protection against an *M. tuberculosis* challenge, a concept that can now be tested in other animal models in a perspective of a putative clinical trial.

## Introduction

*Mycobacterium tuberculosis*, the principal causative agent of human tuberculosis (TB), remains a major global health threat [1] and improved strategies are needed to better control the global TB situation. However, the only currently licensed anti-TB vaccine used at a large scale to vaccinate newborns and infants, mainly in countries with high TB incidence, is represented by the Bacille Calmette and Guerin (BCG) vaccine, an attenuated variant of *Mycobacterium bovis* which has been generated more than 100 years ago [2, 3]. Despite the good protection conferred against severe, disseminated forms of TB in children, the very wide use of BCG has been not been able to stop the global TB epidemic, which is driven by pulmonary TB cases in adolescents and adults that may occur despite BCG vaccination [4, 5]. Among the various factors that could be involved in the limited protection provided by BCG vaccination against pulmonary TB, the absence or solely partial secretion of certain key mycobacterial antigens, as well as the rather weak ability of BCG vaccines to induce long-lasting CD8^+^ T memory responses, might play a role [6][7]. From previous work it is known that BCG lacks a 9.5 kb-sized genomic region, called Region of Difference 1 (RD1) [8][9], which in tubercle bacilli is encoding the ESX-1 type VII secretion system involved in the secretion of key proteins for host-pathogen interaction [10, 11]. To overcome these potential short-comings of BCG, we have previously constructed a recombinant BCG strain, named BCG::ESX-1*^Mmar^* that heterologously expresses the *esx-1* region of *Mycobacterium marinum*, an aquatic mycobacterium that harbors an ESX-1 system similar to *M. tuberculosis* [7]. The introduction of this ESX-1 system into BCG has enhanced host innate immune signaling and induced broader adaptive T-cell responses against ESX-1-secreted proteins compared to parental BCG, while maintaining low virulence in immunocompromised SCID mice, a feature that is different to a related recombinant BCG strain named BCG::RD1 or BCG::ESX-1*^Mtb^* that expresses the ESX-1 system of *M. tuberculosis* and which shows enhanced protective efficacy combined with enhanced virulence in preclinical models [12][13].

To improve anti-TB vaccination efficacy, several possibilities show promise, including (i) development and use of more protective vaccine strains, (ii) adoption of more efficient routes of administration and (iii) employment of heterologous prime-boost vaccination protocols (priming with live-attenuated prophylactic vaccine and boosting once or more with booster vaccine before or after *M. tuberculosis* exposure/infection). Combination of these strategies may avoid the development of active TB disease and prevent late reactivation of the infection ([4][14]).

Among the various routes of administration used for vaccination with different anti-TB vaccine candidates, the mucosal route for immunisation represents a promising choice with multiple advantages over the classical intradermal or subcutaneous routes of immunisation in mouse and NHP animal models [15][16][17]. Since the lungs are the site of *M. tuberculosis* entry and initial infection and also represent the locus for reactivation of the TB disease, the pulmonary mucosal vaccination has been a focus of interest [16]. Indeed, the mucosal immunisation mimics the natural route of *M. tuberculosis* infection and triggers a rapid expansion of T-cell immunity and other innate immune effectors in the airways which may confer improved protection against development of TB disease [18][19][17, 20].

In the present study, we have thus explored aerosol vaccination of mice with BCG::ESX-1*^Mmar^*as well as BCG Pasteur in comparison with subcutaneous vaccination using the same two strains. Our results provide evidence of early key innate cytokine responses with higher frequencies of Th1 effector memory (TEM) cells and significantly greater humoral IgG and IgA responses in the airways of aerosol vaccinated mice compared to the subcutaneous vaccination route. Importantly, only aerosol vaccination was able to induce a substantial resident memory CD4^+^ and CD8^+^ T (TRM) cells in the primary site of *M. tuberculosis* infection leading to significant improved TB protection and lower lung pathology compared to subcutaneous counterparts.

Finally, we show that the BCG::ESX-1*^Mmar^* vaccine candidate displayed a superior T-cell immunity and enhanced anti-mycobacterial protective efficacy in the lungs and spleens when mice were vaccinated via aerosol or subcutaneous routes compared to the parental BCG Pasteur strain. Our data suggest that an aerosol-based mucosal vaccination with BCG::ESX-1*^Mmar^*represents a promising concept worth to be considered for the development of next generation vaccine concepts against *M. tuberculosis*.

## Results

### • Aerosol immunisation of mice with different BCG vaccine strains elicits local and transient inflammatory responses without clinical or tissue pathologies

The main objectives of this work were the evaluation and comparison of host specific immune responses, safety and protective efficacity of BCG::ESX-1*^Mmar^* and parental BCG Pasteur in an aerosol vaccination mouse model relative to the standard subcutaneous vaccination model. To do so, C57BL/6J female mice were vaccinated with either ca. 1 x 10^3^ CFU/mouse via the aerosol route or with 5 x 10^5^ CFU/mouse via the subcutaneous route, as confirmed by CFU counting at day 1 post-immunisation (Figure 1A-B). As expected, neither BCG Pasteur nor BCG::ESX-1*^Mmar^* bacteria were detected in the organs of subcutaneously immunised mice at any time-point post-vaccination, whereas the presence of the vaccine strains was highest in the lungs and spleens of aerosol-immunised mice at four weeks post-immunisation, after which numbers started to decline (Figure 1B). The mycobacterial loads were correlated with host innate pro-inflammatory cytokine responses detected in the airways of the vaccinated mice (Figure 1C). We observed higher IL-1β, IL-6 and TNF-α productions at four weeks post-immunisation and that the BCG::ESX-1*^Mmar^*-immunised mice elicited significantly higher amounts of IL-1β compared to BCG Pasteur counterparts in the early phase following aerosol vaccination (Figure 1C), which is in accordance with the observed *in vitro* ESX-1-dependent enhanced inflammasome activation [7].

**Figure 1.**
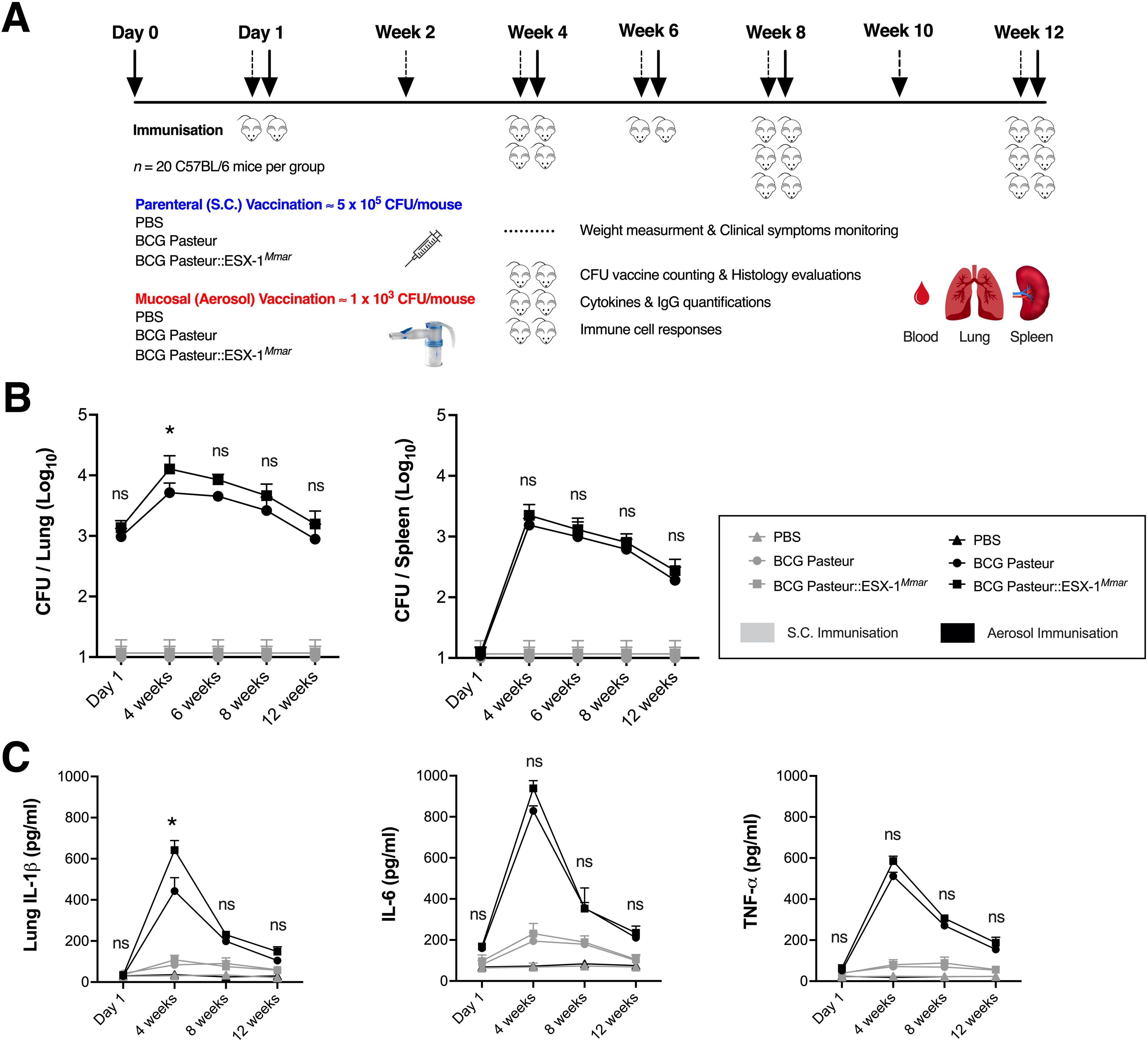
Comparative evaluation model of host immune response of different vaccine strains in mice immunised via different routes. **A.** Protocol adopted in order to evaluate the safety, immunogenicity of the live-attenuated vaccines and their longitudinal interaction with host innate and adaptive immune systems. C57BL/6J female mice (*n* = 20 per group) either left untreated or immunised subcutaneously with 5 x 10^5^ CFU/mouse or via aerosol route with ≈ 1 x 10^3^ CFU/mouse of BCG Pasteur or BCG::ESX-1*^Mmar^* vaccine. At different time-points post-immunisation (solid arrows), vaccine persistence/dissemination, histology and immune responses were evaluated in the lungs, spleens and sera of mice **B.** CFU counting of BCG Pasteur or BCG::ESX-1*^Mmar^*vaccine recovered from the lungs and spleens of vaccinated mice at different time-points post-immunisation (*n* = 2 per group, compiling data of two independent experiments). **C.** Kinetic of pro-inflammatory innate cytokine responses in the lungs of vaccinated mice at different time-points (*n* = 2 per group), as quantified by ELISA. Error bars represent SD. NS = not significant and * = statistically significant with *p*<*0.05*, as determined by Unpaired *t* test with Welch’s correction. The results are representative of at least two independent experiments. Additional data are provided in Suppl. Fig. 1.

Despite the local inflammation observed in the lung of aerosol-immunised mice, practically no significant body weight loss nor any other pronounced clinical symptoms were observed at any time-point following aerosol or subcutaneous vaccination (Suppl. Fig. 1A). In contrast to the lungs of subcutaneously immunised mice, which displayed a normal lung histology at eight weeks following vaccination with BCG Pasteur or BCG::ESX-1*^Mmar^*, histological evaluation of the lungs of aerosol-immunised mice showed important immune cell infiltrations in the perivascular and peribranchial spaces without obvious tissue lesions (Suppl. Fig.1B). We noted qualitatively similar innate and adaptive immune cell infiltration profiles in the inflamed regions of the lungs for BCG Pasteur- and BCG::ESX-1*^Mmar^*-aerosol-immunised mice, which were mainly composed of T lymphocytes (Suppl. Fig. 1C). There were also higher proportions of monocytes/macrophages and neutrophils in the airways of aerosol-vaccinated mice compared to mice vaccinated via the subcutaneous route, as evaluated by microscopy observations and immunohistochemistry on lung sections (Suppl. Fig. 1D).

### • Aerosol immunisation is highly immunogenic and elicits both humoral and cellular T-cell responses in the airways of vaccinated mice

We investigated the impact of vaccination delivery route on the immunogenicity of the vaccine and the magnitude of T cell responses. Our results show that aerosol vaccination with BCG Pasteur or BCG::ESX-1*^Mmar^* induced significantly higher amounts of mycobacteria-specific T-cell IFN-γ responses in the lungs compared to the subcutaneous route at eight weeks post-immunisation (Figure 2A). In addition, BCG::ESX-1*^Mmar^* vaccine triggers robust Th1 responses specific to the major ESX-1-secreted virulence factors and protective immunogens ESAT-6 (EsxA) and CFP-10 (EsxB) (Figure 2A). In contrast to subcutaneous route, only aerosol immunisation was able to elicit notable IL-17A cytokine responses in the airways of BCG Pasteur- and BCG::ESX-1*^Mmar^*-vaccinated mice (Suppl. Fig. 2A).

**Figure 2.**
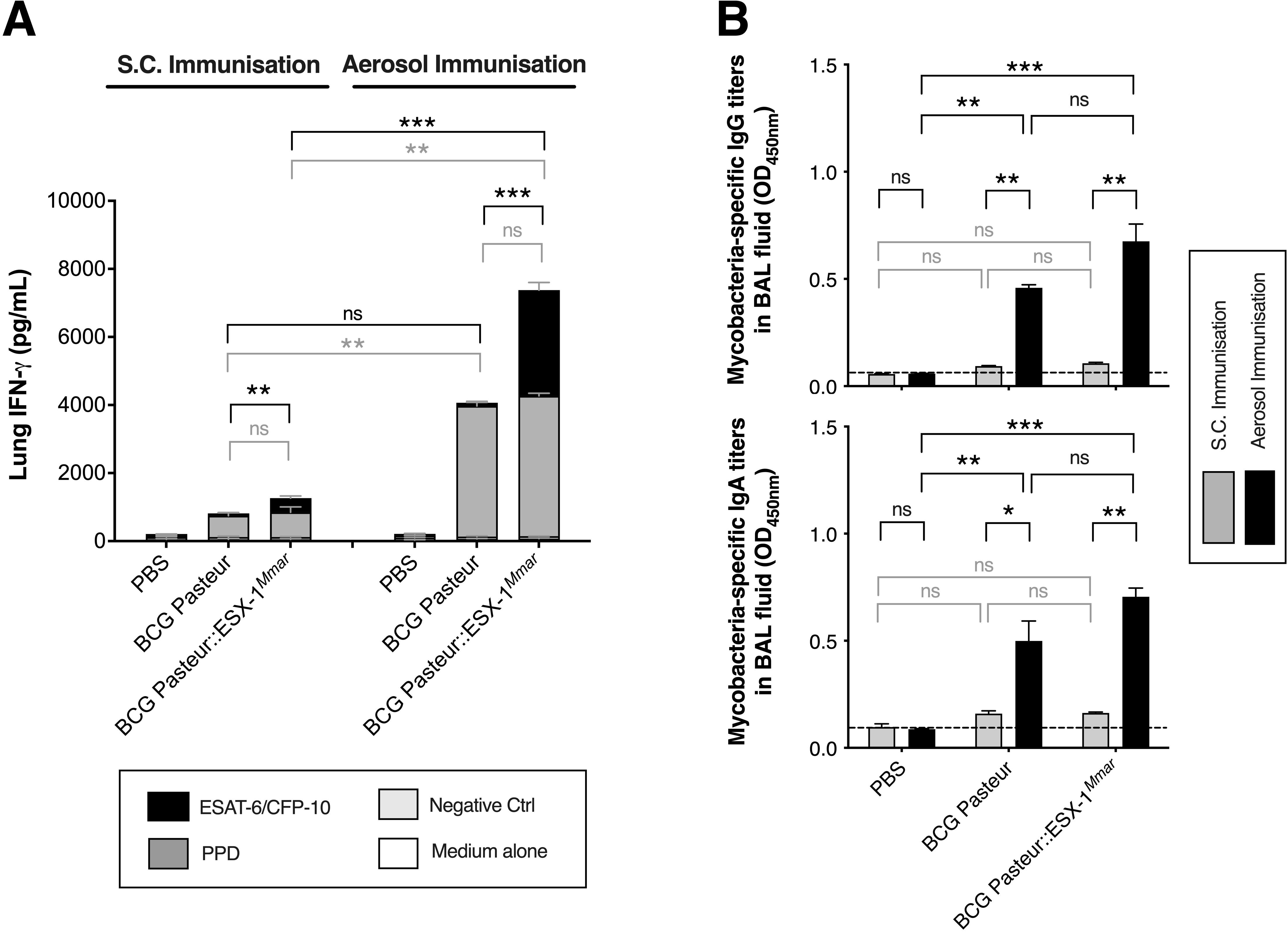
Immunogenicity of tested vaccine strains in mice immunised via subcutaneous or aerosol routes. **A.** T-cell IFN-γ responses in the lungs of BCG Pasteur or BCG::ESX-1*^Mmar^*-vaccinated C57BL/6J mice via subcutaneous or aerosol route, as assessed at eight weeks post-immunisation. Total lung cells of the immunised mice were pooled (*n* = 2 per group) and stimulated *in vitro* with Purified Protein Derivative (PPD), rESAT-6 and rCFP-10 proteins during 72 hours at 37°C. Medium alone and rMalE protein were used as negative controls. The IFN-γ was quantified in the culture supernatant by ELISA. **B.** Humoral immunoglobulin responses in the airways of vaccinated mice at eight weeks post-immunisation (*n* = 2 per group), as evaluated by mycobacterial-specific IgG and IgA titers recovered from the BAL fluid by ELISA. Error bars represent SD. NS = not significant, *, ** and *** = statistically significant with *p<0.05*, *p<0.005* and *p<0.0005*, respectively, as determined by Unpaired *t* test with Welch’s correction. The figures were elaborated by using Prism software. The results are representative of two independent experiments. Additional data are provided in Suppl. Fig. 2-4).

Moreover, we detected high amounts of mycobacteria-specific IgG and IgA responses in the BAL fluid of aerosol-vaccinated mice at eight weeks post-immunisation while subcutaneously immunised mice failed to induce such humoral responses (Figure 2B).

### • Aerosol immunisation triggers higher frequencies of activated CD4^+^ and CD8^+^ T lymphocytes and T effector memory (TEM) cell responses in the lungs compared to subcutaneous route

Induction of robust Th1 effectors subsequent to vaccination is crucial for host defense and correlates with better protection and/or control of *M. tuberculosis* infection [7] [21]. In our model, we detected significant higher frequencies of activated CD4^+^ and CD8^+^ T lymphocytes, recruited into the lung parenchyma of aerosol-immunised mice compared to subcutaneously immunised counterparts following immunisation and also noted increased percentages of such T cell subsets that displayed down-regulation of CD27, CD45RB and CD62L expressions, in the lungs at eight weeks post-immunisation (Suppl. Fig. 3). Their frequency in the airways was lower than observed at four weeks post-immunisation (Suppl. Fig. 4), suggesting a correlation between the mycobacterial burden, host inflammatory responses and recruitment of IFN-γ-producing activated T cells. In addition, a significantly higher number of total CD3^+^ CD4^+^ and CD3^+^ CD8^+^ T lymphocytes were recovered from the lungs of aerosol-immunised mice compared to subcutaneously immunised mice (Suppl. Fig. 1C).

In our model, an increased recruitment/accumulation of both CD4^+^ and CD8^+^ T effector memory cells (TEM defined as CD3^+^ CD28^-^ CD44^high^ CD62L^-^ CD95^+^) in the lungs of aerosol-immunised mice compared to subcutaneous counterparts was observed at eight weeks post-immunisation (Figure 3). These T cells expressed the CXCR3 marker (Figure 3A-B) and showed high intensity of CD44 expression in lung CD4^+^ and CD8^+^ T lymphocytes (Figure 3C), indicating a strong activation and memory function of host Th1 responses subsequent to aerosol vaccination.

**Figure 3.**
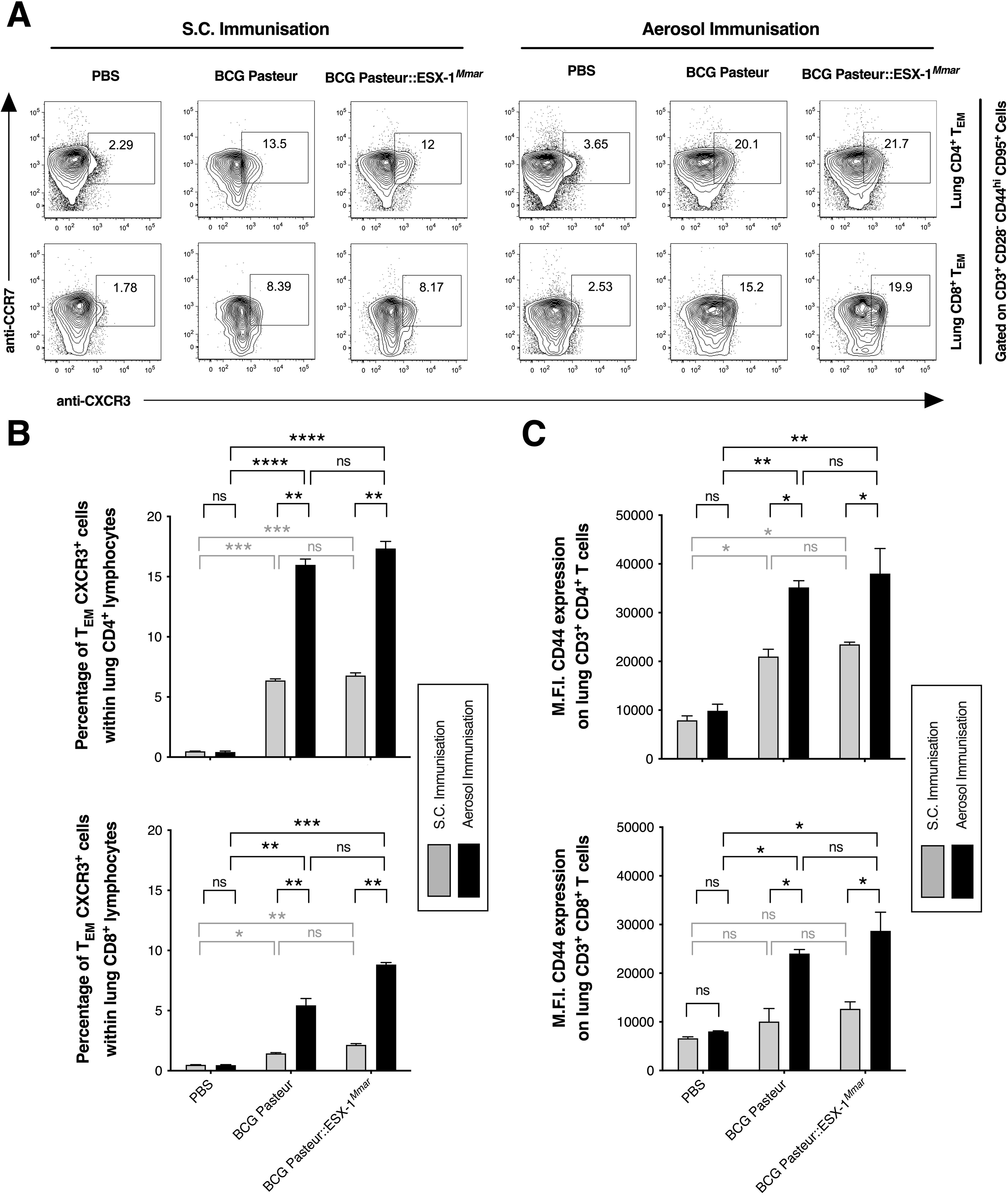
Induction of T effector memory (TEM) cells in the lung of mice immunised with different BCG vaccines via subcutaneous or aerosol route. A-B. T effector memory CD4^+^ and CD8^+^ cells expressing CXCR3 marker recovered from the lungs of BCG Pasteur- or BCG::ESX-1*^Mmar^*-immunised mice via subcutaneous or aerosol route (*n* = 2 per group). Cell populations were gated on CD3^+^ CD28^-^ CD44^high^ CD95^+^ T lymphocytes (**A**). Cytometric plots represent 5% contours with outliers, representative of pool of three mice per group. **B.** Frequency of lung T effector memory CD4^+^ and CD8^+^ (TEM) cells of vaccinated mice among CD4^+^ and CD8^+^ T lymphocyte populations, as evaluated by flow cytometric analyses at eight weeks post-immunisation. **C.** Mean Fluorescence Intensities (M.F.I.) of CD44 expression on total lung CD3^+^ CD4^+^ and CD3^+^ CD8^+^ T cells recovered from the same vaccinated mice (**A-B**). Error bars represent SD. NS = not significant, *, **, ***, and **** = statistically significant with *p<0.05, p<0.005, p<0.001* and *p<0.0001*, respectively, as determined by Unpaired *t* test with Welch’s correction. At least 500,000 events per sample were acquired. The obtained data were analyzed using FlowJo software and figures were elaborated by using Prism software. The results are representative of two independent experiments. Additional data are provided in Suppl. Fig. 5.

### • Only aerosol immunisation was able to elicit substantial T resident memory (TRM) cell responses in the airways of vaccinated mice

Previous mouse models have suggested that mucosal vaccination induces lung-associated T resident memory (TRM) cells that protect against TB disease [22, 23]. Hence, we evaluated the capacity of different routes of immunisation with BCG Pasteur and BCG::ESX-1*^Mmar^* vaccines to induce notable long-lived TRM cells in our mouse model. We found that only aerosol vaccination elicited substantial CD69^+^ CD103^+^ TRM lymphocytes, containing both CD4^+^ and CD8^+^ T cell compartments at eight weeks post-immunisation, while subcutaneous immunisation failed to induce such response (Figure 4). These cell populations are defined as CD3^+^ CD45^+^ CD44^high^ CD62L^-^ CD69^+^ CD103^+^ cells (Figure 4A) in mice. Importantly, the BCG::ESX-1*^Mmar^* vaccine generated significantly higher percentages and absolute numbers of lung CD4^+^ and CD8^+^ TRM cell subsets in the airways of aerosol-immunised mice compared to BCG Pasteur (Figure 4B-C).

**Figure 4.**
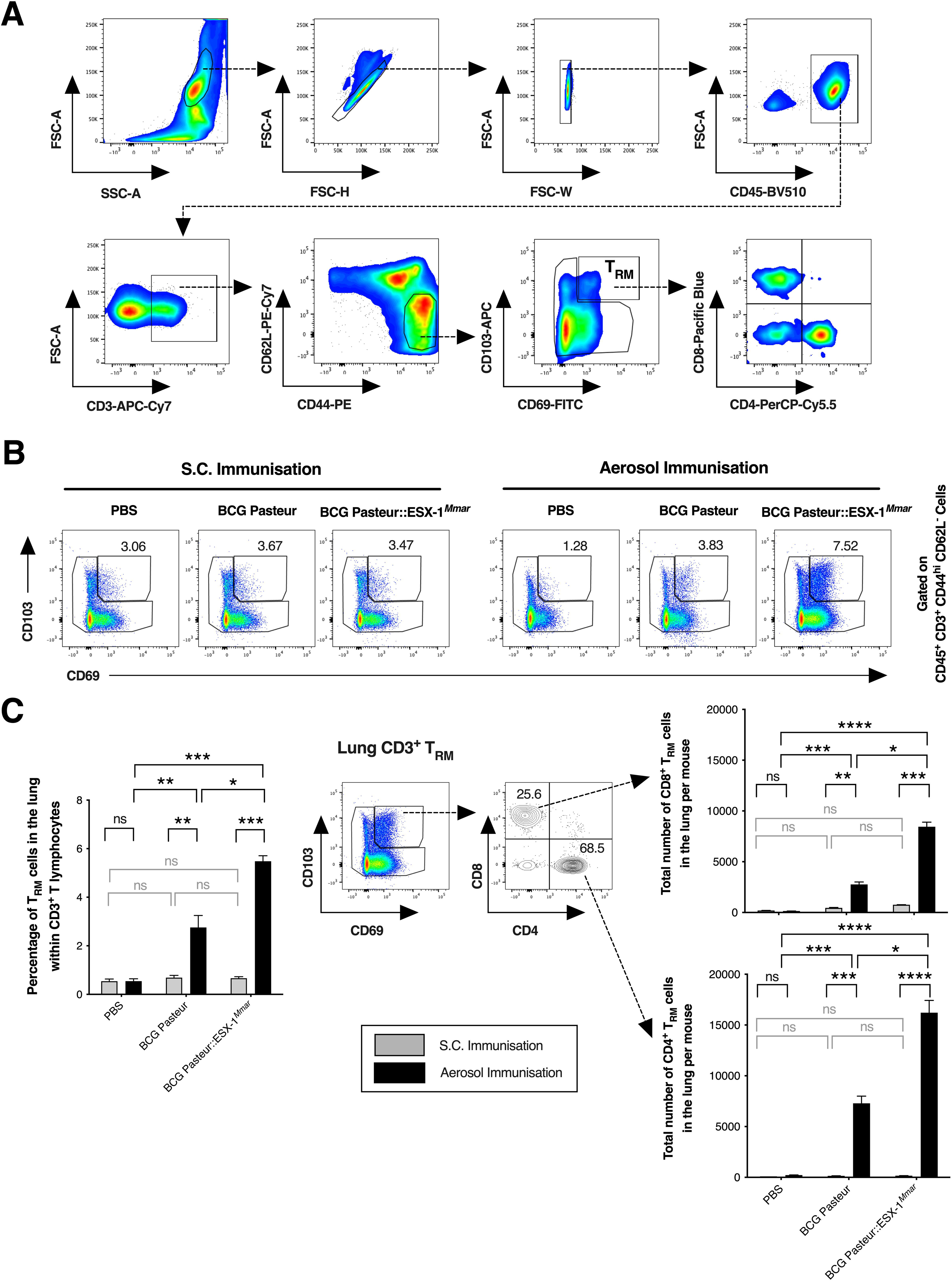
Induction of T resident memory (TRM) cells in the lungs of mice immunised with different BCG vaccines via subcutaneous or aerosol routes. **A.** Gating strategy and flow cytometric analyses adopted to identify both lung CD4^+^ and CD8^+^ T resident memory (TRM) cell populations. **B.** Frequency of CD69^+^ CD103^+^ TRM cells recovered from the lung parenchyma of mice vaccinated with BCG Pasteur or BCG::ESX-1*^Mmar^* via subcutaneous or aerosol route (*n* = 2 per group), as evaluated by flow cytometric analyses at eight weeks post-immunisation. **C.** Percentages and absolute numbers of lung CD4^+^ and CD8^+^ TRM cells per mouse. Error bars represent SD. NS = not significant, *, **, ***, and **** = statistically significant with *p<0.05, p<0.005, p<0.001* and *p<0.0001*, respectively, as determined by Unpaired *t* test with Welch’s correction. At least 500,000 events per sample were acquired, representative of pool of three mice per group. The obtained data were analyzed using FlowJo software and figures were elaborated by using Prism software. The results are representative of two independent experiments.

### Aerosol immunisation induces potent polyfunctional Th1 effectors in the spleens and systemic IgG responses similar to those detected after subcutaneous immunisation

In addition to vaccine-induced T-cell immunity in the airways, we investigated the capacity of aerosol immunization to induce Th1 effectors in the secondary lymphoid organs. This exercise showed comparable amounts of mycobacteria-specific T-cell IFN-γ responses in the spleens of aerosol-immunised and subcutaneously immunised mice at eight weeks post-immunisation. Mice vaccinated with BCG::ESX-1*^Mmar^* displayed such responses also against the major ESX-1 antigens ESAT-6 and CFP-10, while this was not the case for BCG Pasteur vaccinated mice, due to the absence of these antigens from parental BCG [7, 24] (Figure 5A). Moreover, we observed strong CD4^+^ and CD8^+^ TEM cell responses in the spleens of these mice (Suppl. Fig. 5A), although at a lower frequency in aerosol vaccinated mice relative to the ones vaccinated by the subcutaneous route (Suppl. Fig. 5B-C). In-depth characterization of the functional Th1 cell subsets and their differentiation status by intra-cellular cytokine staining (ICS) showed that aerosol and subcutaneous immunisations with BCG Pasteur or BCG::ESX-1*^Mmar^*induced globally comparable Th1 effectors with similar compositions of IL-2, TNF-α and IFN-γ cytokine-producing CD4^+^ T subsets in the spleens of vaccinated mice at eight weeks post-immunisation upon *in vitro* stimulation with PPD (Figure 5B). The responses were dominated by terminally differentiated single-positive TNF-α+ and IFN-γ+ CD4^+^ T cells, followed by double-positive TNF-α+ IL-2^+^ CD4^+^ T cells. We noted that single-positive IFN-γ+ CD4^+^ T cells were slightly higher in the spleens of subcutaneously immunised mice while single-positive TNF-α+ CD4^+^ T cells were lower than in aerosol-immunised mice. Furthermore, comparable CCR6 and CXCR3 expression percentages on polyfunctional IL-2^+^ TNF-α^+^ IFN-γ^+^ cytokine-producing CD4^+^ T cell subsets were detected in the spleens of all vaccinated mice (Figure 5B).

**Figure 5.**
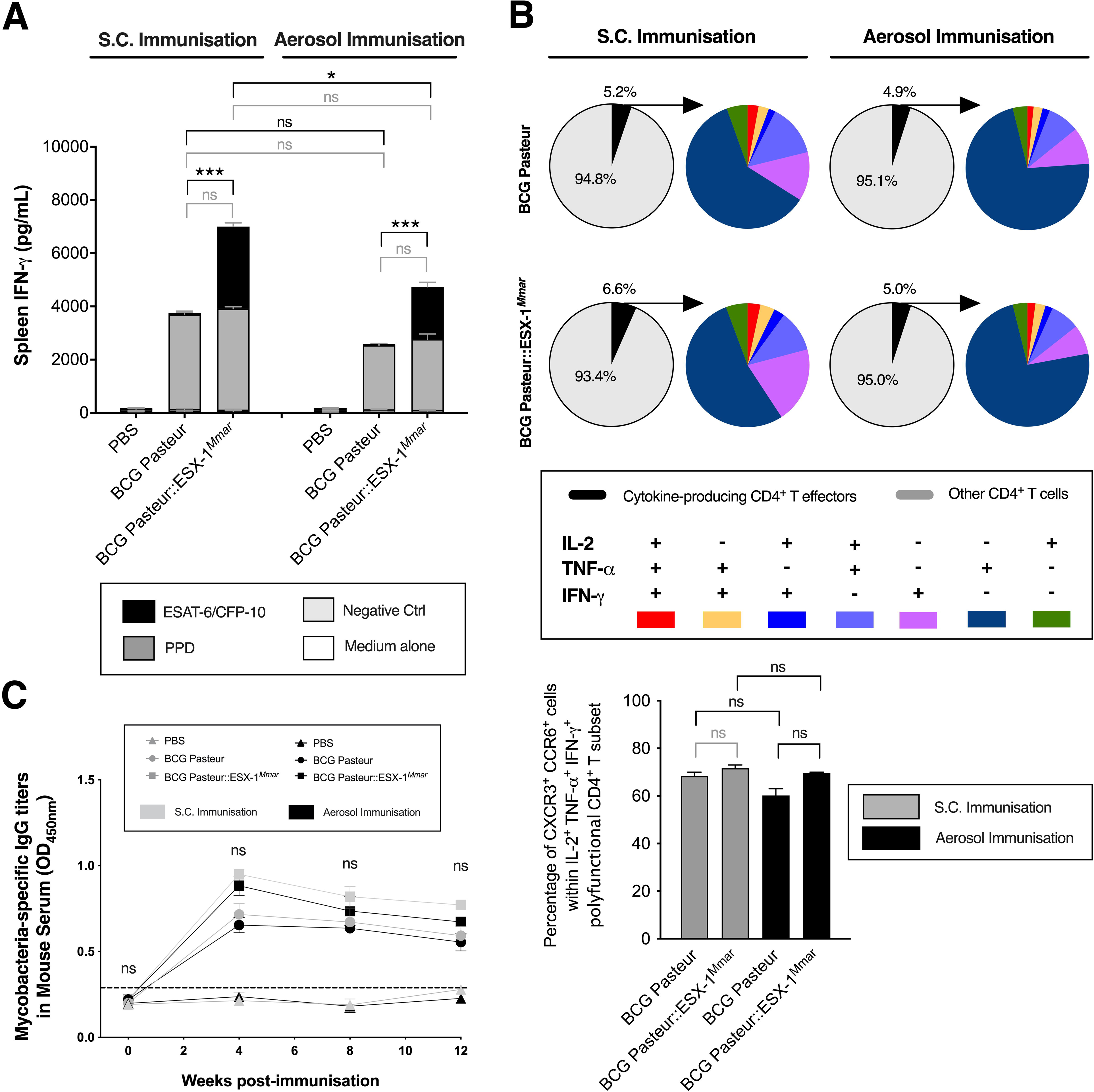
Induction of polyfunctional Th1 cell responses in the spleens and IgG responses in the sera of mice vaccinated with different BCG vaccines via subcutaneous or aerosol route. **A.** T-cell IFN-γ responses in the spleens of BCG Pasteur or BCG::ESX-1*^Mmar^*-vaccinated mice via subcutaneous or aerosol route, as assessed at eight weeks post-immunisation. Pool of total splenocytes (*n* = 2 per group) were stimulated *in vitro* with mycobacterial antigens and the IFN-γ was quantified in the culture supernatant by ELISA as described in Figure 2A. **B.** Frequency of Th1 cytokine-producing CD4^+^ T subsets in the spleens of vaccinated mice at eight weeks post-immunisation. Total splenocytes of vaccinated mice (*n* = 2 per group) were stimulated *in vitro* with 10 μg/ml of PPD for 24 hours prior to ICS staining (IL-2, TNF-α and IFN-γ) and flow cytometric analyses. At least 1,000,000 events per sample were acquired for flow cytometric analyses. The obtained data were analyzed using FlowJo software and figures were elaborated by using Prism software. The results are representative of two independent experiments. **C.** Mycobacteria-specific IgG titers in the sera of vaccinated mice at different time-points post-immunisation (*n* = 2 per group), as evaluated by ELISA. Error bars represent SD. NS = not significant, * and *** = statistically significant with *p<0.05* and *p<0.001*, respectively, as determined by Unpaired *t* test with Welch’s correction. The figures were elaborated by using Prism software. Additional data are provided in Suppl. Fig. 6-7.

In addition, we detected comparable mycobacteria-specific cytokine-producing CD8^+^ T effectors in the spleens of mice independent of the vaccine route at twelve weeks post-immunisation (Suppl. Fig. 6A-B). A higher percentages of TNF-α+ and IFN-γ^+^ bi-functional CD8^+^ T lymphocytes were induced by BCG::ESX-1*^Mmar^* compared to BCG Pasteur, probably due to broader responses to ESX-1 secreted antigens and/or induction of cytosolic immune signalling for the ESX-1-proficient strain compared to BCG Pasteur, leading to enhanced MHC-I-restricted epitope presentation [7][25][26]. In contrast, the composition of Th1 cytokine-producing CD4^+^ T effectors in the spleens of mice twelve weeks post-immunisation was comparable in the BCG::ESX-1*^Mmar^* and the BCG Pasteur groups (Suppl. Fig. 6C), similar to what has also been observed at eight weeks post-immunisation. Also, the total numbers of both CD4^+^ and CD8^+^ T lymphocytes in the spleens of vaccinated mice were comparable between both vaccination routes at twelve weeks post-immunisation (Suppl. Fig. 6D).

As regards humoral responses, it was noticed that aerosol and subcutaneous vaccinations elicited notable and long-term systemic humoral responses comparable at all time-points following vaccination, as judged by PPD-specific IgG titers quantified in the sera of vaccinated mice (Figure 5C). This finding suggests that aerosol vaccination with live-attenuated vaccines is a highly immunogenic route able to elicit not only greater local immune responses compared to the parenteral route, but also comparable systemic responses.

### Aerosol vaccination of mice significantly improved TB protection and showed lower lung pathology compared to parenteral vaccination route

Finally, we wondered if the enhanced lung innate immunity and higher mycobacteria-specific Th1 activated/effector memory responses and CD4^+^ and CD8^+^ TRM cells induced by aerosol immunisation correlated with improved protection in mice against an *M. tuberculosis* challenge. The experiments showed that mice immunised with BCG Pasteur or BCG::ESX-1*^Mmar^*via aerosol route displayed significant improved TB protection in the lungs compared to subcutaneously immunised mice, as judged by reduction of mycobacterial loads in the lungs at four weeks post-infection (Figure 6B), as well as in term of lower lung pathology (Figure 7). Indeed, fewer tissue lesions and cell infiltrations were found the lungs of BCG Pasteur- and BCG::ESX-1*^Mmar^*-vaccinated mice via the aerosol route compared to subcutaneously vaccinated groups at four weeks post-infection (Figure 7). In contrast, no statistically significant differences in CFU counts were noted between aerosol and subcutaneous vaccination routes in the spleens of mice at four weeks post-infection (Figure 6C). For these mouse experiments we also noticed that vaccination with BCG::ESX-1*^Mmar^* was more protective than vaccination with BCG Pasteur for both immunisation routes, when using CFU counts as a readout (Figure 6B-C), as well as in depth histopathological evaluations on some key parameters such as percentages of lung damaged and consolidated areas, and inflammation seriousness (Figure 7A-B).

**Figure 6.**
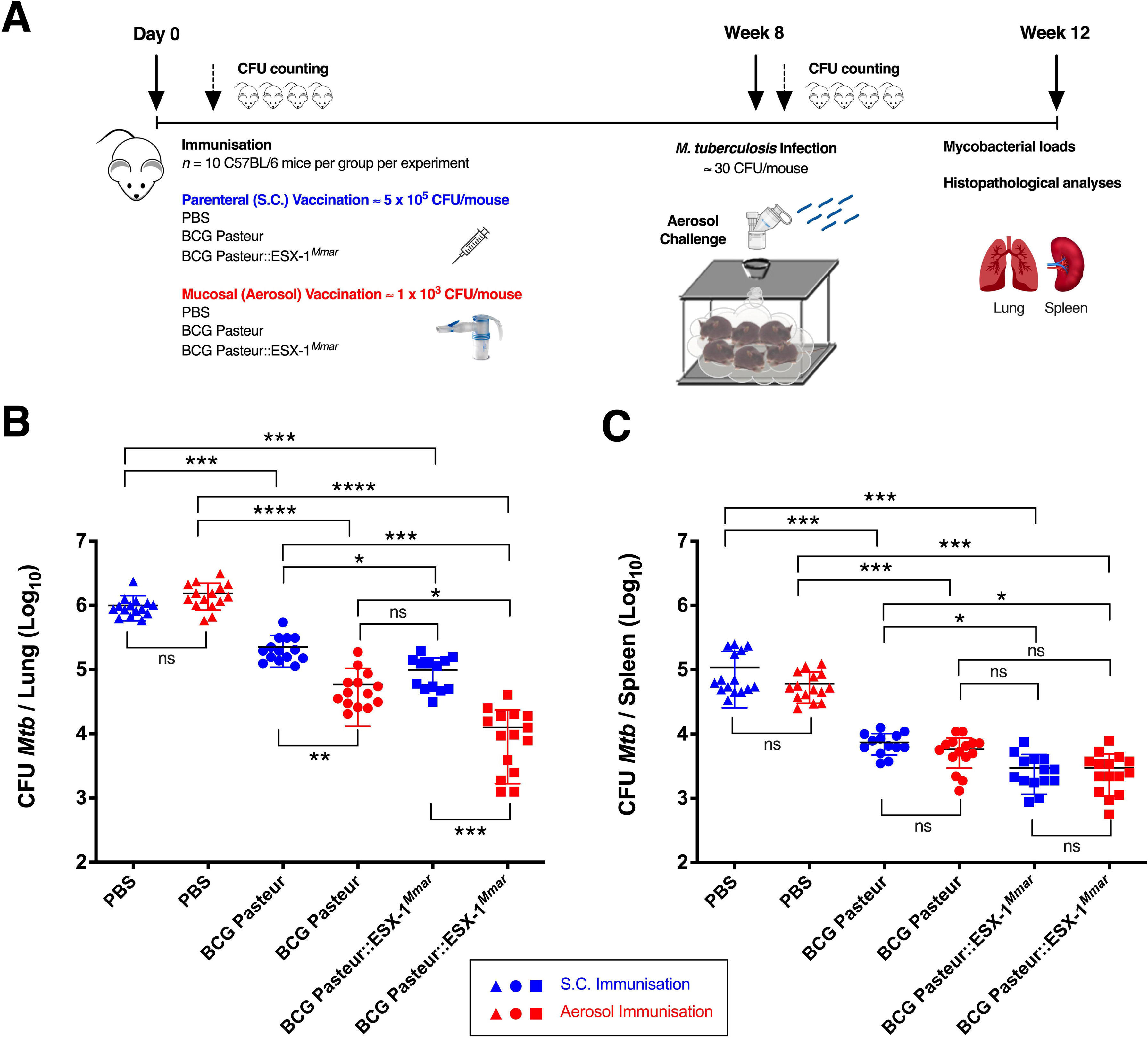
Comparative protection efficacy of different vaccine strains in mice immunised via subcutaneous or aerosol route against an *M. tuberculosis* challenge. A. Scheme to evaluate vaccine protection efficacy against infection with low dose of *M. tuberculosis* H37Rv strain via aerosol route. C57BL/6J mice (*n* = 10 per group, compiling data of two independent experiments) either left untreated or immunised subcutaneously with 5 x 10^5^ CFU/mouse or via aerosol route with ≈ 1 x 10^3^ CFU/mouse of BCG Pasteur or BCG::ESX-1*^Mmar^* vaccine. Four weeks post-infection, the mycobacterial loads in the lungs and spleens of individual mice were quantified by CFU counting and lung histopathological evaluations were performed. **B-C.** C57BL/6J mice were immunised with BCG Pasteur or BCG::ESX-1*^Mmar^* vaccine strains via subcutaneous (blue) or aerosol (red) route. Mice were infected with ≈ 30 CFU/mouse of *M. tuberculosis* H37RV WT strain via aerosol route at eight weeks post-immunisation. The mycobacterial load in the lung (**B**) and spleen (**C**) of individual mice were determined by CFU counting at four weeks post-infection. NS = not significant, *, **, ***, **** = statistically significant with *p*<*0.05*, *p*<*0.005*, *p*<*0.0005* and *p*<*0.0001*, respectively, as determined by Brown-Forsythe & Welch ANOVA tests for Multiple Comparisons with Dunnett’s T3 Corrections. The results are accumulation of two independent experiments and figures were elaborated by using Prism software.

**Figure 7.**
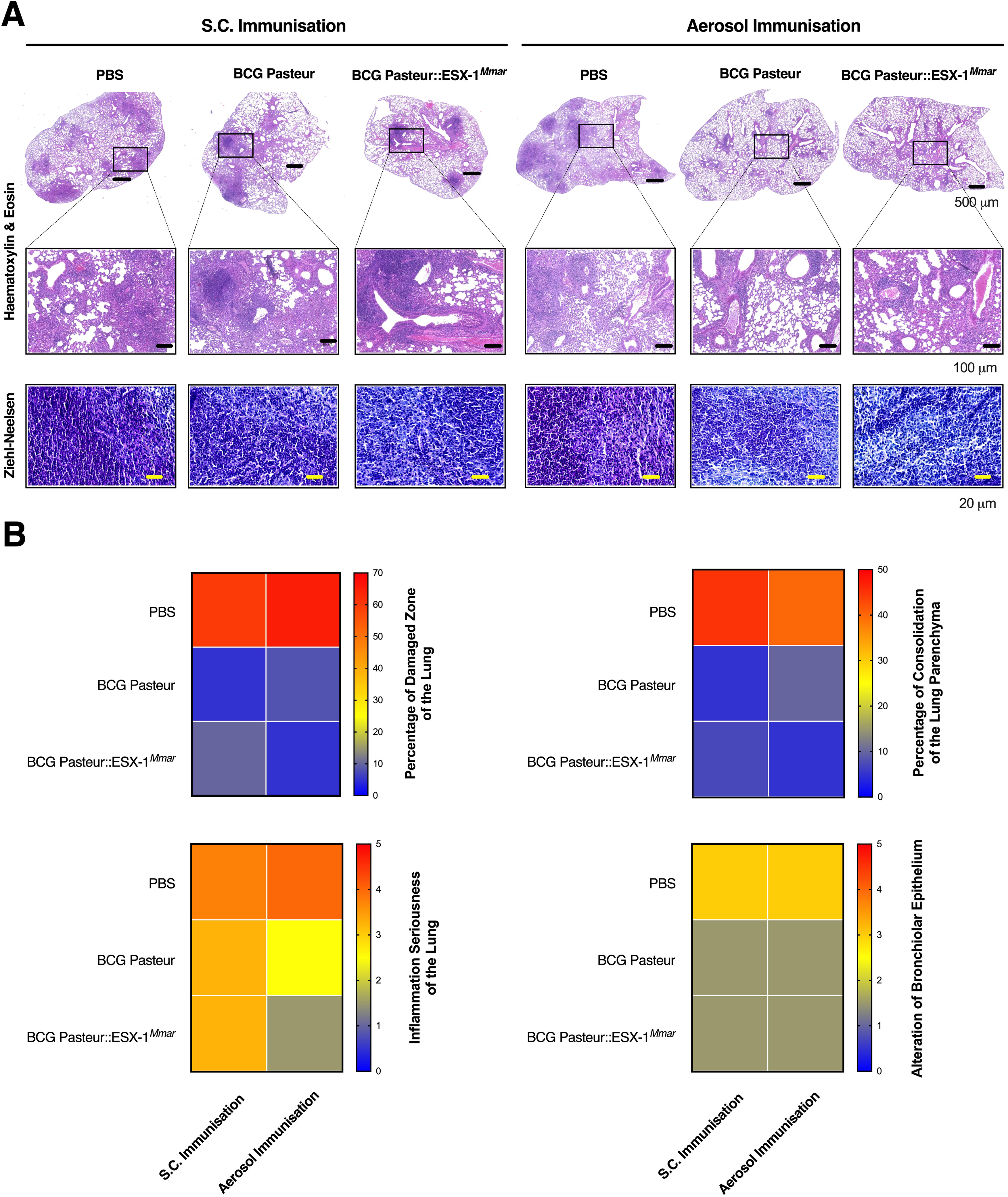
Histological evaluations of the lungs of mice vaccinated with different BCG vaccines via subcutaneous or aerosol route and infected with *M. tuberculosis*. A. Histopathological analyses on lung sections (left lobe) of *M. tuberculosis*-infected mice showing lesions and cell infiltrations, as evaluated by hematoxylin and eosin (H&E) staining method and acid-fast bacilli were observed by Ziehl-Neelsen staining method at four weeks post-infection (**A**). Scale bars represent 500 μm (upper row), 100 μm (middle row) and 20 μm (lower row). **B.** Comparative lung histopathological evaluations and median scoring notes, based on different histological parameters, attributed to each group of mice. Five-scale severity grade was adopted for histopathological scoring; 1: minimal, 2: mild, 3: moderate, 4: marked and 5: severe. The results are representative of two independent experiments and figures were generated and elaborated by using Zen and Prism softwares, respectively.

Together, these results obtained with our murine model highlight the potential advantages of the aerosol vaccination route compared to the classical route of administration of live-attenuated anti-TB vaccines and also indicate that the BCG::ESX-1*^Mmar^* vaccine represents an improved candidate for such a revised strategy of vaccination against *M. tuberculosis*.

## Discussion

To achieve the WHO goal of enhanced control and potential eradication of TB worldwide, more effective vaccines and new anti-TB drugs are needed. An optimized vaccination strategy against TB should primarily provide a better protection of adolescents and adults, both in terms of Protection of Infection (POI) and Protection of Disease (POD), as individuals from these age-sections are often - despite BCG vaccination in their early childhood - not efficiently protected against pulmonary forms of TB disease, which represent the most efficient transmission routes for spreading *M. tuberculosis* to new hosts, keeping the TB infection cycle going.

Development of improved vaccination strategies against infection and/or TB disease in adolescents and adults, resulting either from primo-infection, reactivation and/or re-infection, thus represents a key objective for preclinical and clinical research. One of the premises for generating improved vaccine candidates is to increase the immunogenicity relative to the standard BCG vaccines, whereby live-attenuated vaccine candidates such as VPM1002 [27][28] and/or MTBVAC [29][30] are currently evaluated in clinical trials.

In a similar attempt, we have recently generated a recombinant BCG Pasteur strain complemented with an integrating cosmid that carries the *esx-1* region of *M. marinum* [7]. The heterologous expression of the *M. marinum* ESX-1 type VII secretion system in this vaccine candidate impacted different aspects of host-pathogen interaction, including the induction of phagosomal rupture, the access to the host cytosol and enhanced immune signaling [7]. These aspects are all important parameters during mycobacterial infection, with profound influence on host innate and adaptive immune responses [31][32][33][34][7][21][25]. Indeed, the BCG::ESX-1*^Mmar^* strain was able to modulate the host innate immune response via phagosomal rupture-associated induction of type I interferon (IFN) responses that were transmitted through the cGAS/STING pathway, as well as enhanced inflammasome activity, resulting in higher IL-1β release and higher proportions of CD8^+^ T cell effectors against mycobacterial antigens and potent poly-functional CD4^+^ Th1 cells specific to ESX-1-secreted antigens ESAT-6, CFP-10 and EspC [7][25]. We hypothesize that these changes are probably the main reason for the superior TB protection observed in mice vaccinated with BCG::ESX-1*^Mmar^* compared to BCG Pasteur and BCG Danish vaccine strains in independent murine infection models that used the H37Rv reference strain, but also hypervirulent *M. tuberculosis* strains belonging to Beijing and Haarlem strain families as challenge strains [7].Given these encouraging previous results we also included the BCG::ESX-1*^Mmar^*strain in our current study, which mainly focused on exploring different vaccination routes.

Indeed, in the current study, we compared aerosol-mediated vaccination versus standard subcutaneous vaccination in C57BL6J mice using BCG Pasteur and BCG::ESX-1*^Mmar^* as live-attenuated vaccine strains, evaluating their immunogenicity and space-temporal interactions with the host. As reported in detail in the results section, aerosol vaccination with BCG Pasteur and BCG::ESX-1*^Mmar^* was highly immunogenic, inducing strong innate and adaptive activated/effector immune cell responses at the airway mucosal (local) level (Figures 1-3) as well as systemic poly-functional Th1 cytokine-producing cell responses that were comparable between the two vaccination routes (Figure 5 and Suppl. Fig. 5-6).

Interestingly, only aerosol vaccination induced substantial long-lived non-circulating CD4^+^ and CD8^+^ T resident memory (TRM) cells in the lung tissue of immunised mice while subcutaneous vaccination failed to do so (Figure 4). These results are in good agreement with examples from the literature, indicating that TRM cells play numerous immune functions and have tissue-specific and migration property-specific differentiation programs [35][36, 37]. As such it was shown that mouse CD4^+^ and CD8^+^ TRM cells share immunosurveillance strategies in the lung tissue after acute viral infection and that CD4^+^ TRM reactivation plays a key role in the immune recall by triggering chemokine responses and amplifying immune cell activation [37]. Zens et al., 2016 reported that long-term protective CD4^+^ and virus-specific CD8^+^ TRM cells were only generated in the mouse airways subsequent to intranasal vaccination with live-attenuated influenza vaccine, similar in phenotype to those generated by influenza virus infection, but not following vaccination with injectable inactivated influenza vaccine, a finding that strongly suggests that the induction of these responses is highly dependent on the nature of vaccine and the administration route [38][35].

In additional to cell-mediated immunity [39], B cells and humoral antibody responses in the mucosa of vaccinated mice may efficiently contribute to host defense and the control of *M. tuberculosis* infection [40][41]. Whereas elevated titers of IgG and IgM responses were present in the BAL fluid of aerosol-vaccinated mice, and not for subcutaneously vaccinated mice (Figure 2B), serum antibody titers displayed strong and persistent serum IgG responses in both vaccination routes (Figure 5C).

Nemeth et al., 2020 showed in a mouse model, which used a contained and persistent yet non-pathogenic infection with *M. tuberculosis*, that rapid and durable reduction of TB disease burden was achieved in these mice when they were re-exposed through an aerosol challenge with the same *M. tuberculosis* strain. This experimental model, which mimics to some extend a latent TB infection showed that pre-exposure to low dose *M. tuberculosis* can be beneficial for the host, protecting to some degree against a subsequent aerosol challenge with higher doses. The observed protection is associated with elevated activation of alveolar macrophages and accelerated recruitment of *M. tuberculosis*-specific T cells to the lung parenchyma, resulting in reduced disease upon re-exposure to *M. tuberculosis* [42]. Moreover, another recent study revealed that a low-level persistent infection of mice with *Plasmodium chabaudi* generated protective immunity by impacting CD4^+^ T cell subsets, promoting in particular both IFN-γ^+^ and TNF^+^ double positive and terminally differentiated IFN-γ^+^ T effectors [43]. These results support the hypothesis of a potentially beneficial role of persistent microbial infections at a non-pathogenic level, which may create an immunologic condition that is reminiscent to the one generated by the use of live-attenuated vaccine strains.

Our study also demonstrated that mice vaccinated with BCG Pasteur or BCG::ESX-1*^Mmar^* via the aerosol route were significantly better protected against a challenge with virulent *M. tuberculosis* compared to mice vaccinated via the subcutaneous route, as shown by reduced mycobacterial loads (Figure 6B) and superior lung histology scores with lower inflammatory lesions/disease seriousness (Figure 7). Interestingly, comparable *M. tuberculosis* CFU counts were observed in the spleens of these aerosol-vaccinated mice relative to subcutaneously vaccinated mice, even with an administrated vaccine dose that was ≈ 500-fold lower in the aerosol group. These findings suggest that similar levels of CD4^+^ and CD8^+^ TEM responses were induced in the spleens by both vaccination routes despite different doses (Figure 5 and Suppl. Fig. 5-6), leading to a comparable protection level in this organ (Figure 6C). Concerning the lungs, we hypothesize that the superior protection seen in aerosol-vaccinated mice relative to subcutaneously vaccinated mice is likely due to an enhanced quantity and quality of sustainable T-cell immunity in this site, involving TEM, TRM and humoral IgA and IgG responses.

An increased protection conferred by mucosal vaccination relative to subcutaneous vaccination was also observed in several previous studies using murine infection models. For example, it has been shown that intranasal administration of BCG in mice induced superior protection in the lungs at early time-points as well as long elevated protective splenic responses with higher frequencies of CD4^+^ and CD8^+^ T cells expressing IFN-γ [44]. A greater protective efficacy following intranasal BCG administration was also observed in TB-susceptible DBA/2 mice, with higher *M. tuberculosis*-specific Th1 and Th17 immune responses, as well as IgA concentration in lungs compared to subcutaneous vaccination [45]. The mucosal (both intranasal and intratracheal) BCG vaccination in mice also generated T effector memory and T resident memory cells in the lung [22] and was able to induce potent lung tissue resident PD-1^+^ KLRG1^-^ CD4^+^ T cells [46] that confer enhanced pulmonary protection against *M. tuberculosis*. In addition, Mata et al., 2021, showed that pulmonary but not subcutaneous administration of BCG induces lung-resident macrophage activation in mice, even after vaccine clearance [47]. It was also reported that this activation of alveolar macrophages was mainly mediated by CD4^+^ T cells, providing not only a long-term TB protection but also conferring heterologous protection against *Streptococcus pneumoniae* infection, suggesting that BCG mucosal vaccination drives potent *in vivo* trained innate memory-like responses. Such an additional protective effect of trained immunity might also be induced by mucosal vaccination with BCG Pasteur and BCG::ESX-1*^Mmar^* vaccine, and might contribute to the improved protection observed in our model. Indeed, immunohistochemistry analyses revealed a significant higher number of F4/80^+^ activated monocytes/macrophages in the lung parenchyma of aerosol-vaccinated mice compared to subcutaneously vaccinated mice (Suppl. Fig. 1D).

In conclusion, our experiments in mice revealed that both the vaccine route as well as the vaccine type influenced the degree of protection from a challenge with *M. tuberculosis*. Benefits were seen for aerosol vaccination compared to subcutaneous vaccination in terms of CFU reduction and histological preservation of the lung tissue. Moreover, superior sustained CD4^+^ and CD8^+^ TEM and TRM responses were induced by the vaccination with BCG::ESX-1*^Mmar^* compared to the BCG Pasteur parental strain, as observed at the time of challenge with *M. tuberculosis* (Figures 3-4). This finding suggests that T-cell immunity mediated by ESX-1-secreted antigens efficiently contributes to enhanced TB protection in the lungs of mice vaccinated with BCG::ESX-1*^Mmar^* compared to BCG Pasteur for both tested vaccination routes (Figure 6-7). As such, our study confirms that BCG::ESX-1*^Mmar^* does represent an interesting new vaccine candidate and further suggests that it is a promising asset for conventional, subcutaneous as well as alternative, aerosol-based vaccination, worth to be considered for further preclinical and eventual clinical development.

## Experimental Procedures (Materials and Methods)

### Mycobacterial strains

*M. bovis* BCG Pasteur was grown in Dubos broth medium (Becton Dickinson) supplemented with Albumine, Dextrose and Catalase (ADC, Becton Dickinson) at 37°C and BCG::ESX-1*^Mmar^*was grown in the same medium to which hygromycin 50 µg/ml was added. Mycobacterial concentrations were determined by OD600nm measurement and CFU counting on Middlebrook 7H11 solid Agar medium (Becton Dickinson) complemented with Oleic acid, Albumine, Dextrose and Catalase (OADC, Becton Dickinson) after 3-4 weeks of culture at 37°C.

*M. tuberculosis* H37Rv WT strain used for mouse infection experiments was prepared from titrated frozen stocks kept at -80°C that were originally prepared from growing cultures in Dubos broth complemented with ADC at 37°C. Mycobacterial suspensions were washed twice with PBS and left to rest for 20 minutes before being collected as single-cell suspension. CFU were counted on Middlebrook 7H11 solid Agar medium complemented with OADC after 3-4 weeks of incubation at 37°C.

### Animal studies and ethical statement

All animal-involving studies were conducted in agreement with the European and French guidelines (EC Directive 2010/63/UE and French Law 2013-118 issued on 1 February 2013). These experiments were approved by the Institut Pasteur safety committee (protocol 11.245) and by the relevant Ethics Committee (Comité d’éthique en experimentation animale 89) and by the French Ministry for Higher Education and Research (dap 180023, APAFIS #15409– 2018060717283847 v1 and dap220021, APAFIS #37011–2022042811231522 v1), respectively.

### Protection assay, organ preparations and CFU counting

Six-to seven-week-old female C57BL/6J mice (Janvier) were used for protection studies. Mice were immunised subcutaneously with ≈ 5 x 10^5^ CFU/mouse of different BCG vaccine strains (100 μl of volume given at the base of the tail) or via aerosol route generated from suspensions containing 2.5 x 10^7^ CFU/ml of BCG vaccine strains in order to reach an inhaled dose of ≈ 1 x 103 CFU of vaccine per mouse lung. Aerosol vaccination route was generated from mycobacterial suspension using clinically proven and commercial nebulizer (Pari LC Sprint) attached to mice-containing isolation chamber in L3 protection animal facility. Mice were placed in custom-made individual tubes in order to maximize homogeneous dose distribution. The received dose in the lungs was evaluated by CFU counting of lung homogeneates from 2-3 mice up to 24 hours post-immunisation. Organs from the different groups of mice were used for CFU counting and histological analyses and others were used for immunological evaluations at different time-points post-immunisation.

Mice were infected via aerosol route with a low dose of *M. tuberculosis* H37Rv strain (≈ 30 CFU/mouse), as evaluated by CFU counting in the lung up to 24 hours post-infection. Four weeks later, mice were culled and lungs and spleens harvested for CFU counting and histopathological analyses. Individual organs were homogenized using a MillMixer organ homogenizer (Qiagen) and serial 5-fold dilutions were plated on Middlebrook 7H11 solid Agar medium complemented with 10% OADC, and/or containing Ampicillin (50 μg/ml), PANTA mixture (Polymyxin B, Amphotericin B, Nalidixic acid, Trimethoprim and Azlocillin, Becton Dickinson) or no antibiotics. CFU were counted after 3-4 weeks of incubation at 37°C.

### Immune cell preparations, staining and flow cytometry

Adaptive immune cells from the lungs and spleens of immunised mice were prepared as previously described [48, 49]. Briefly, lungs were washed with PBS and digested by treatment with 400 U/ml type-IV collagenase and DNase-I (Roche) for 30 minutes at 37°C. Single-cell suspensions were prepared by use of a GentleMacs (Miltenyi) and by passage through 100-μm diameter filters (Cell Strainer, BD Falcon). Lung and spleen cell suspensions were enriched in live lymphocytes T and B by using a density gradient centrifugation Ficoll medium (Lympholyte-M, Cedarlane) according to the manufacturer’s protocol. The obtained lymphocyte layers were washed twice with PBS and incubated with appropriate dilutions of mAb containing different combinations of anti-CD3-APC-Cy7, anti-CD4-PerCP-Cy5.5, anti-CD8-Pacific Blue, anti-CD27-APC, anti-CD28-FITC, anti-CD44-PE, anti-CD45-BV510, anti-CD45RB-FITC, anti-CD62L-PE-Cy7, anti-CD69-FITC, anti-CD95 (Fas)-APC, anti-CD103-APC, anti-CD183 (CXCR3)-BV510, anti-CD196 (CCR6)-PE-Cy7 and anti-CD197 (CCR7)-PE-Cy7, prepared in FACS buffer (PBS containing 3% fetal bovine serum and 0.1% NaN3) during 30 minutes at 10°C sheltered from light. The stained cells were washed twice with FACS buffer and then fixed with an appropriate volume of 4% paraformaldehyde overnight at 10°C prior to sample acquisition. At least 500,000 events per sample, depends on experiment, were acquired by using LSR Fortessa flow cytometer system and BD FACSDiva software (BD Bioscience). The obtained data were analyzed using FlowJo software (Treestar) and graphs were performed by Prism 10 software (GraphPad Prism).

### T-cell stimulation and intracellular cytokine staining (ICS) assay

Single-cell suspensions from spleens of immunised mice were obtained by tissue dissociation, homogenization and passage through 100 μm-pore filter. Pooled cells (*n* = 3 mice /group) were cultured at 7.5 x 10^6^ cells/well in flat-bottom 12-well plates (TPP) in the presence of 1 μg/ml anti-CD28 (clone 37.51) and 1 μg/ml of anti-CD49d (clone 9C10-MFR4.B) mAbs (BD Pharmingen) and stimulated with 10 μg/ml of Purified Protein Derivative (PPD) during 24 hours at 37°C et 5% CO2 in RPMI 1640 medium GlutaMAX (Gibco), complemented with 10% fetal bovine serum (Gibco) followed by 4-5 hours of incubation with protein transport inhibitor containing Brefeldin A (Golgi Plug, BD Pharmingen), according to constructer’s instructions. Cells were then harvested, washed twice with FACS buffer and incubated for 10 minutes at 10°C with FcγII/III receptor blocking anti-CD16/CD32 (BD 2.4G2 clone) mAb. Cells were then incubated for 30 minutes with appropriate dilutions of anti-CD3ε-APC-eFluor780, anti-CD4-BUV737 and anti-CD8α-Pacific Blue mAbs (BD Pharmingen) at 10°C and sheltered from light. The stained cells were washed twice with FACS buffer, permeabilized by use of Cytofix/Cytoperm kit (BD Pharmingen). Cells were then washed twice with PermWash buffer (BD Pharmingen) and incubated with appropriate dilutions of PerCP-Cyanine5.5-anti-IL-2 (clone JES6-5H4, eBioscience), PE-anti-TNF-α (clone 554419, BD Pharmingen), and Alexa Fluor647-anti-IFN-γ (clone XMG1.2, eBioscience) mAbs during 30 minutes at 10°C. Appropriate staining with control Ig isotypes was performed in parallel. Cells were subsequently washed with PermWash buffer and then with FACS buffer before fixation with 4% paraformaldehyde for 2-4 hours at 10°C. At least 1,000,000 events per sample, were acquired by using LSR Fortessa flow cytometer system and BD FACSDiva software (BD Bioscience). The obtained data were analyzed using FlowJo software (Treestar) and graphs were performed by Prism 10 software (GraphPad Prism).

### T-cell assay and ELISA

Pool of splenocytes and total lung homogenates of immunised mice (*n* = 2-3 mice per group) were cultured in flat-bottom 24-well plates (TPP) at 5 x 10^6^ cells per well in RPMI 1640 medium GlutaMAX (Gibco), complemented with 10% fetal bovine serum (Gibco), 1% of MEM non-essential amino acids (Invitrogen, Life Technologies), 5 x 10^-5^ M β-mercaptoethanol (Invitrogen, Life Technologies), 100 U/ml penicillin and 100 µg/ml streptomycin (Sigma-Aldrich) in the presence of 5 µg/ml of PPD (Creative Diagnostics), or a mixture of 2 µg/ml rESAT-6 (Abcam) and 2 µg/ml rCFP-10 (RayBiotech) proteins during 72 hours at 37°C and 5% CO2. Medium alone and 4 µg/ml of rMalE protein were used as negative controls.

Cytokine productions in culture supernatants were quantified by ELISA. Nunc 96-well Maxisorp plate (Thermo-Fisher) were used and mAbs specific to IFN-γ (clone AN-18 for coating and clone R4-6A2 for detection) or IL-17A (clone TC11-18H10 for coating and clone TC11-8H4.1 for detection) were from BD Pharmingen. Innate cytokine quantifications in the organs of vaccinated mice were performed as described [48, 49].

Total and mycobacteria-specific Immunoglobulin IgG and IgA titers were quantified in the sera and bronchoalveolar lavage (BAL) fluids of vaccinated mice by ELISA. HRP goat anti-mouse IgG (BioLegend) and goat-anti-mouse IgA-HRP (SouthernBiotech) were used for quantification in diluted BAL fluids. Nunc 96-well Maxisorp plate (Thermo-Fisher) were used, and optical density was measured at 450 nm by FLUOstar Omega microplate reader (BMG LABTECH). Graphs were elaborated using Prism 10 software (GraphPad Prism).

### Histology, immunohistochemical studies and histopathological scoring

Samples of respective lung lobes from immunised mice were fixed in 10% neutral buffered formalin for 72 hours, transferred to 70% ethanol and prepared for histopathological evaluations and scoring. Organs were embedded in paraffin for 4 μm microtome sectioning and stained with hematoxylin and eosin (H&E) and/or acid-fast bacilli Ziehl-Neelsen staining methods according to standard procedures. Monoclonal antibodies specific to F4/80 (Cell Signaling, D2S9R), Ly6B.2 (bio rad, MCA771G), B220 (BD Pharmingen, 550286), CD4 (Cell Signaling, D7D2Z), CD8 (Cell Signaling, D4W2Z) surface markers were used for immunohistochemical (IHC) studies in order to identify different immune cell populations in the lungs of mice.

Microscopic changes were qualitatively described and when applicable, scored semi-quantitatively, by histopathologists blinded to the conditions, using distribution qualifiers (i.e., focal, multifocal, locally extensive or diffuse), and a five-scale severity grade, i.e., 1: minimal, 2: mild, 3: moderate, 4: marked and 5: severe.

Analyses were performed at the Histology Platform of the Institut Pasteur and stained slides were evaluated with ZEISS Axio Scan.Z1 Digital Slide Scanner and Zen software version 3.9 (Zeiss).

### Statistical analyses

Graphs and statistical analyses were performed and analyzed using Prism 10 software (GraphPad Prism). The Unpaired *t* test with Welch’s correction and Brown-Forsythe & Welch ANOVA tests for multiple comparisons with Dunnett’s T3 Corrections were employed depending on the data to determine the significance values between the different groups.

## Supporting Information

**Suppl. Fig. 1. Animal well-being and histological evaluations of the lungs of mice immunised with different BCG vaccines via subcutaneous or aerosol route. A.** Body-weight measurement and clinical symptom follow-up evaluations of C57BL/6J mice (*n* = 4 per group) that were either left untreated, immunised subcutaneously with 5 x 10^5^ CFU/mouse or vaccinated via the aerosol route with ≈ 1 x 10^3^ CFU/mouse of BCG Pasteur or BCG::ESX-1*^Mmar^* vaccine. Three-scale severity grades were adopted for mouse clinical scoring; 1: low, 2: moderate, 3: severe. **B.** Histological analyses on lung sections of individual vaccinated mice, evaluated by hematoxylin and eosin (H&E) staining method at eight weeks post-immunisation. Scale bars represent 500 μm (upper row) and 100 μm (lower row). **C.** Total numbers of CD3^+^ CD4^+^ and CD3^+^ CD8^+^ T cells recovered from the lungs of immunised mice at eight weeks post-immunisation, as determined as total numbers of cells in the Ficoll-treated lung fractions, multiplied by the percentages of CD4^+^ or CD8^+^ cells, respectively, as assessed by flow cytometry. **D.** Immunohistochemistry evaluations on lung sections of vaccinated mice detecting macrophage and neutrophil cell infiltrations at eight weeks post-immunisation. Error bars represent SD. NS = not significant, * and ** = statistically significant with *p<0.05* and *p<0.005*, respectively, as determined by Unpaired *t* test with Welch’s correction. The figures were elaborated by using Prism software. The results are representative of two independent experiments.

**Suppl. Fig. 2. IL-17A cytokine responses in the lungs of mice immunised with different BCG vaccines via subcutaneous or aerosol route. A.** Quantification of IL-17A in the lungs of C57BL/6J mice (*n* = 2 per group) vaccinated via subcutaneous or aerosol route as assessed at four weeks post-immunisation. Total lung cells of the immunised mice were stimulated *in vitro* with Purified Protein Derivative (PPD) during 72 hours at 37°C. Medium alone and rMalE protein were used as negative controls. The IL-17A amount was quantified in the culture supernatant by ELISA. **B.** Total numbers of CD3^+^ CD4^+^ and CD3^+^ CD8^+^ T cells recovered from the lungs of immunised mice at four weeks post-immunisation, as determined as total numbers of cells in the Ficoll-treated lung fractions, multiplied by the percentages of CD4^+^ or CD8^+^ cells, respectively, as assessed by flow cytometry. Error bars represent SD. NS = not significant, *, ** and *** = statistically significant with *p<0.05*, *p<0.005* and *p<0.001*, respectively, as determined by Unpaired *t* test with Welch’s correction. The figures were elaborated by using Prism software. The results are representative of two independent experiments.

**Suppl. Fig. 3. Profile of adaptive immune cells in the lungs of mice immunised with different BCG vaccines via subcutaneous or aerosol route at eight weeks post-immunisation.** Expression of T-cell activation/recruitment markers on the surface of lung CD3^+^ CD4^+^ (**A**) and CD3^+^ CD8^+^ (**B**) T lymphocytes, recovered from BCG Pasteur- or BCG::ESX-1*^Mmar^*-immunised C57BL/6J mice via subcutaneous or aerosol route, as evaluated by flow cytometric analyses at eight weeks post-immunisation. At least 500,000 events per sample were acquired. Cytometric plots represent 5% contours with outliers, representative of pool of two mice per group. The obtained data were analyzed using FlowJo software and figures were elaborated by using Prism software. The results are representative of two independent experiments.

**Suppl. Fig. 4. Profile of adaptive immune cells in the lungs of mice immunised with different BCG vaccines via subcutaneous or aerosol route at four weeks post-immunisation.** Expression of T-cell activation/recruitment markers on the surface of CD3^+^ CD4^+^ (**A**) and CD3^+^ CD8^+^ (**B**) T lymphocytes, recovered from BCG Pasteur- or BCG::ESX-1*^Mmar^*-immunised C57BL/6J mice via subcutaneous or aerosol route, as evaluated by flow cytometric analyses at four weeks post-immunisation. At least 500,000 events per sample were acquired. Cytometric plots represent 5% contours with outliers, representative of pool of two mice per group. The obtained data were analyzed using FlowJo software and figures were elaborated by using Prism software. The results are representative of two independent experiments.

**Suppl. Fig. 5. Induction of Th1 effector memory (TEM) cells in the spleens of mice immunised with different BCG vaccines via subcutaneous or aerosol route. A-B.** T effector memory CD4^+^ and CD8^+^ cells expressing CXCR3 marker recovered from the spleens of the same C57BL/6J vaccinated mice in Figure 3 at eight weeks post-immunisation. Cell populations were gated on CD3^+^ CD28^-^ CD44^high^ CD95^+^ T lymphocytes (**A**). Cytometric plots represent 5% contours with outliers, representative of pool of two mice per group. **B.** Frequency of T effector memory CD4^+^ and CD8^+^ cells (TEM) of vaccinated mice among spleen CD4^+^ and CD8^+^ T lymphocyte populations, as evaluated by flow cytometric analyses. **C.** Mean Fluorescence Intensities (M.F.I.) of CD44 expression on total spleen CD3^+^ CD4^+^ and CD3^+^ CD8^+^ T cells, as evaluated by flow cytometric analyses. Error bars represent SD. NS = not significant, *, ** and *** = statistically significant with *p*<*0.05*, *p*<*0.005* and *p*<*0.001*, respectively, as determined by Unpaired *t* test with Welch’s correction. At least 500,000 events per sample were acquired. The obtained data were analyzed using FlowJo software and figures were elaborated by using Prism software. The results are representative of two independent experiments.

**Suppl. Fig. 6. Induction of mycobacteria-specific long-lasting CD8^+^ T cells and polyfunctional Th1 response in the spleens of mice immunised with different BCG vaccines via subcutaneous or aerosol route. A-B.** TNF-α and IFN-γ producing CD8^+^ T lymphocytes by ICS applied on total splenocytes of mice at twelve weeks post-immunization. Pool of splenocytes (*n* = 2 per group) were stimulated *in vitro* with 10 μg/ml of PPD for 24 hours or left untreated prior to staining with cytokine-specific mAbs (**A**) or control immunoglobulin (Ig) isotypes (**B**). At least 500,000 events per sample were acquired for flow cytometric analyses. Cytometric plots are representative of pool of two mice per group. **C.** Frequency of Th1 cytokine-producing cells within spleen CD4^+^ T effectors recovered from vaccinated mice at twelve weeks post-immunisation. Total splenocytes of vaccinated mice (*n* = 2 per group) were stimulated *in vitro* with 10 μg/ml of PPD for 24 hours prior to ICS staining (IL-2, TNF-α and IFN-γ) and flow cytometric analyses. At least 500,000 events per sample were acquired for flow cytometric analyses. **D.** Total numbers of CD3^+^ CD4^+^ and CD3^+^ CD8^+^ T cells recovered from the spleens of vaccinated immunised mice at twelve weeks post-immunisation, as determined as total numbers of cells in the Ficoll-treated lung fractions, multiplied by the percentages of CD4^+^ or CD8^+^ cells, respectively, as assessed by flow cytometry. Error bars represent SD. NS = not significant, as determined by Unpaired *t* test with Welch’s correction. At least 500,000 events per sample were acquired for flow cytometric analyses. The obtained data were analyzed using FlowJo software and figures were elaborated by using Prism software.

**Suppl. Fig. 7. Gating strategy and cytometric analyses adopted to identify different subsets of Th1 cytokine-producing T effectors in the spleens of vaccinated mice. A-B**. Pool of total splenocytes of BCG Pasteur-immunised mice (*n* = 2 per group) stimulated *in vitro* with 10 μg/ml of PPD for 24 hours prior to ICS staining (**A**). The negative controls for ICS assay performed to dissect mycobacteria-specific CD4^+^ T cell responses in the spleens of the same vaccinated mice without stimulation prior to staining with cytokine-specific mAbs (**B**) or with control Ig isotypes. At least 1,000,000 events per sample were acquired for flow cytometric analyses. Cytometric plots represent 5% contours with outliers, representative of pool of two mice per group. The obtained data were analyzed using FlowJo software and figures were elaborated by using Prism software. The results are representative of two independent experiments.

## Acknowledgments

We thank Daria Bottai from the University of Pisa for fruitful discussions and advice.

We also thank the members of the Institut Pasteur A3 animal facility, in particular Mathilde Dubot, Rachid Chennouf and Karim Sebastien for expert animal care. We are also grateful to the team of the Imaging platform of the Institut Pasteur, in particular Pierre-Henri Commere, for help and support.

## Author Contributions

**F.S.** conceptualized, designed and performed the experiments, analyzed the data, generated the figures, wrote and edited the manuscript.

**W.F.** prepared relevant cultures of vaccine strains, performed molecular verifications of the vaccine strains and performed *in vivo* experiments.

**A.P.** performed *in vivo* experiments and managed animal facility logistics.

**C.T.** provided additional *in vitro* analyses.

**M.T.** & **D.H.** performed the histological preparation and staining of the samples, analyzed the data.

**R.B.** conceptualized and designed experiments, supervised the methodology and the results, wrote, edited and submitted the manuscript and acquired the funding.

