## Supplementary figures and images for "Improved Immune Responses and Tuberculosis Protection by Aerosol Vaccination with recombinant BCG expressing ESX-1 from *Mycobacterium marinum*"

### Suppl Fig 1

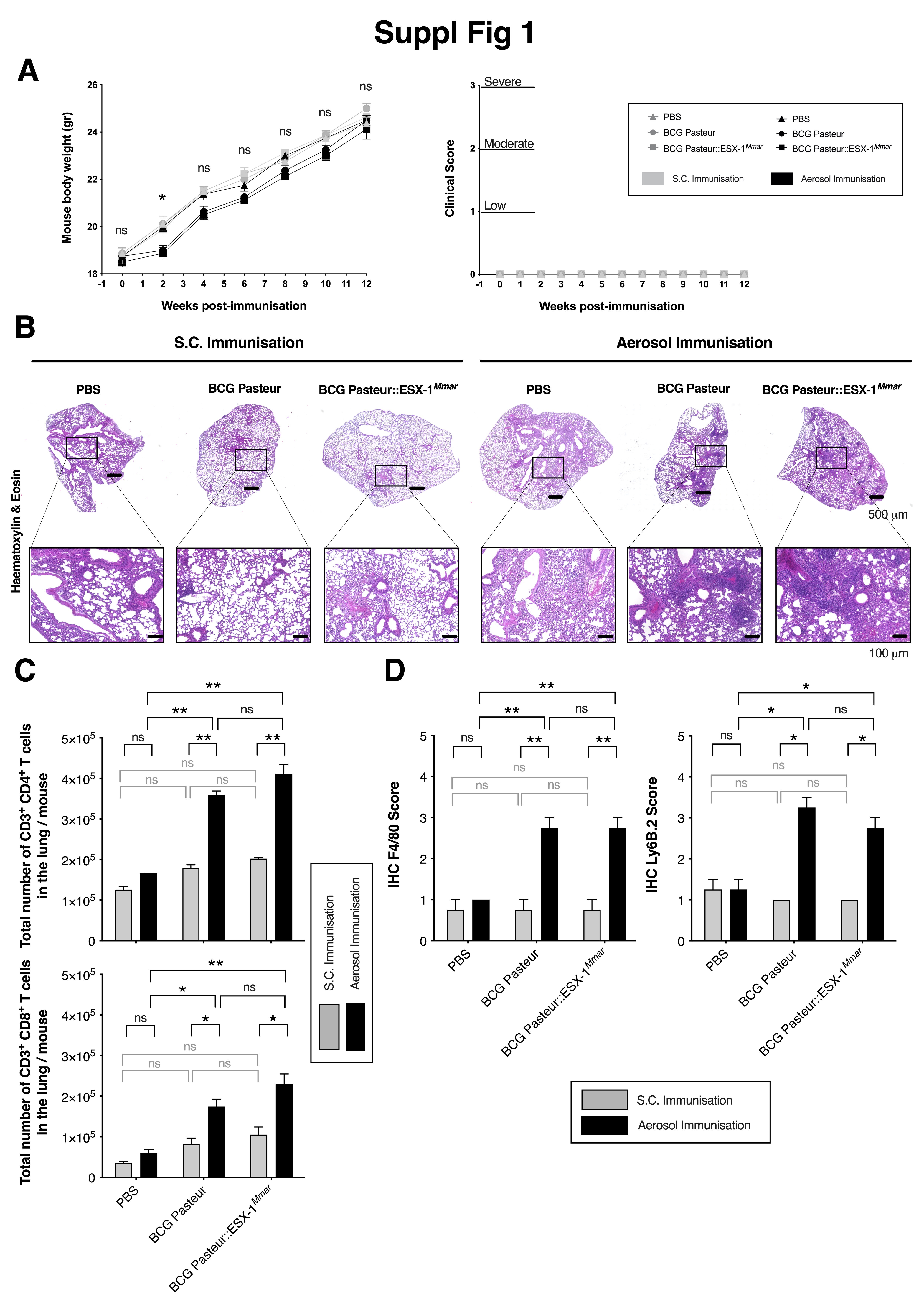

### Suppl Fig 2

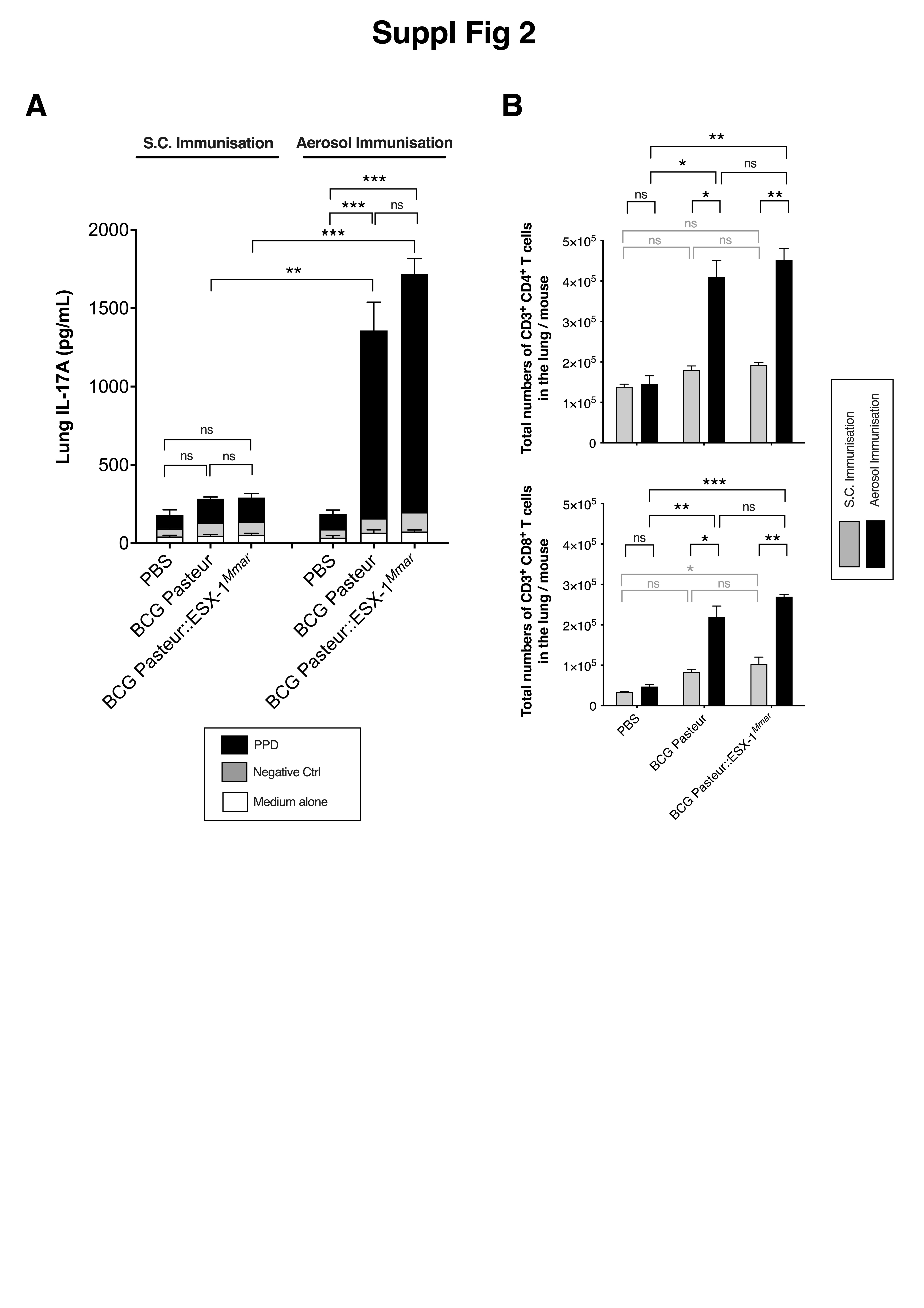

### Suppl Fig 3

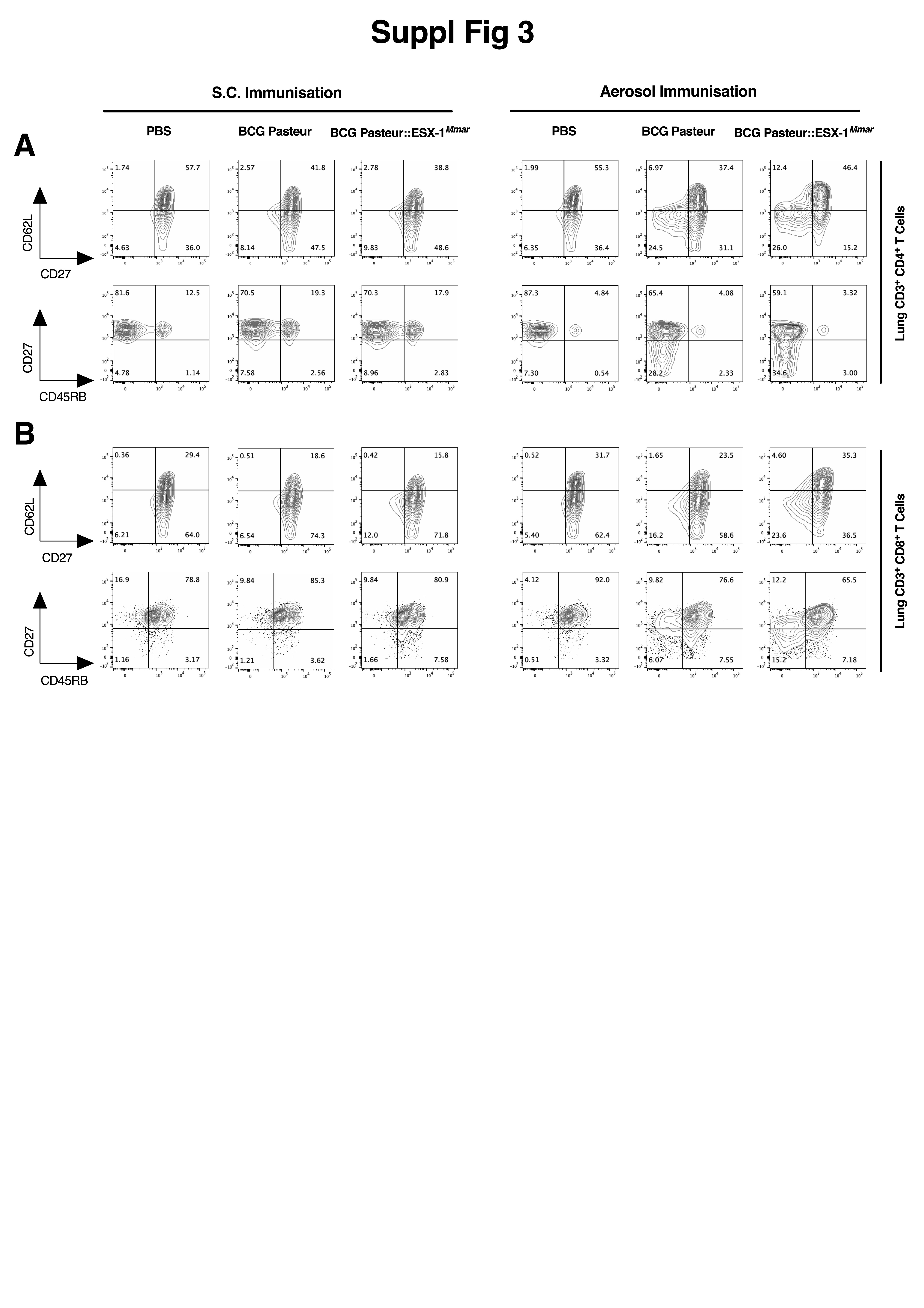

### Suppl Fig 4

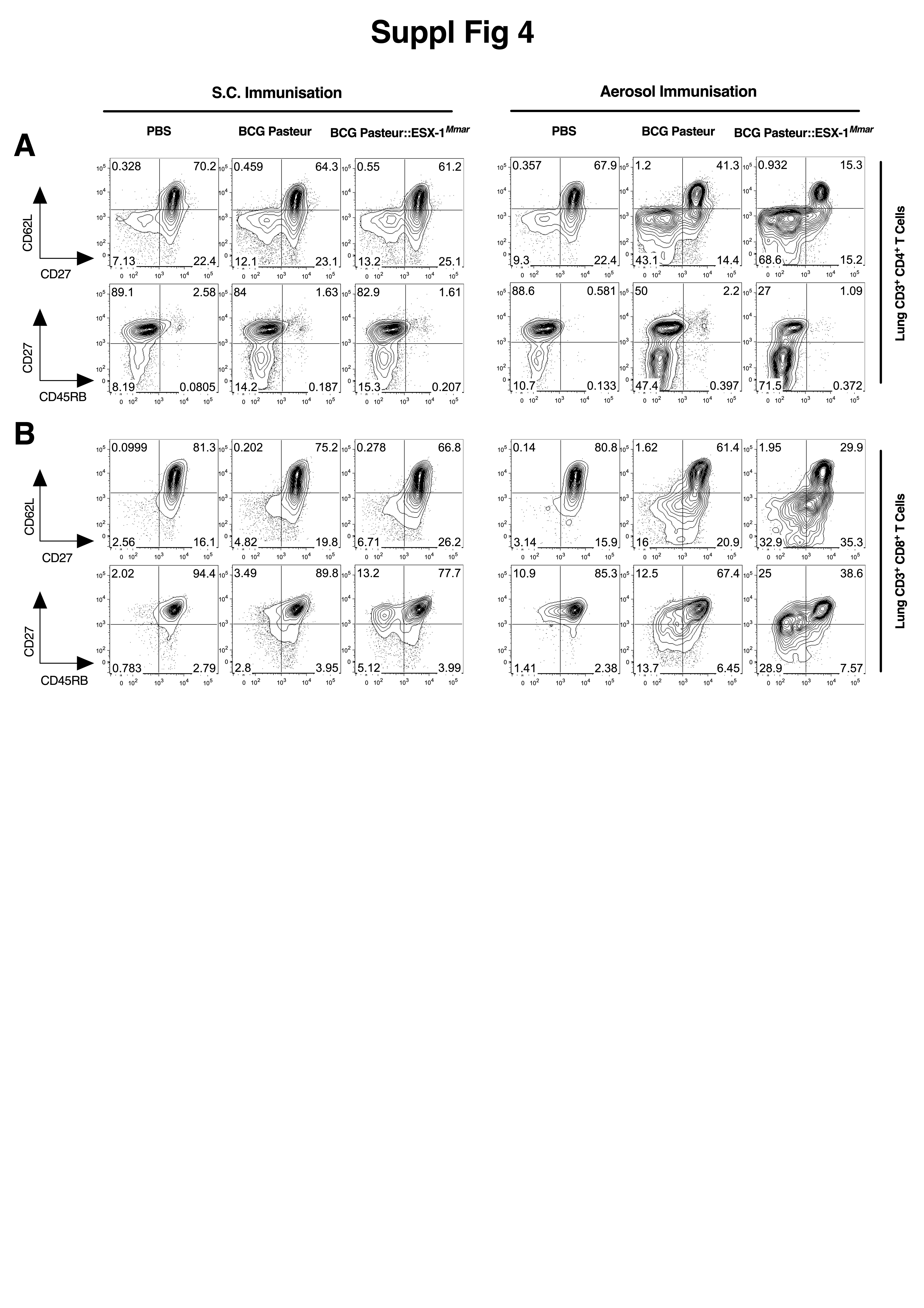

### Suppl Fig 5

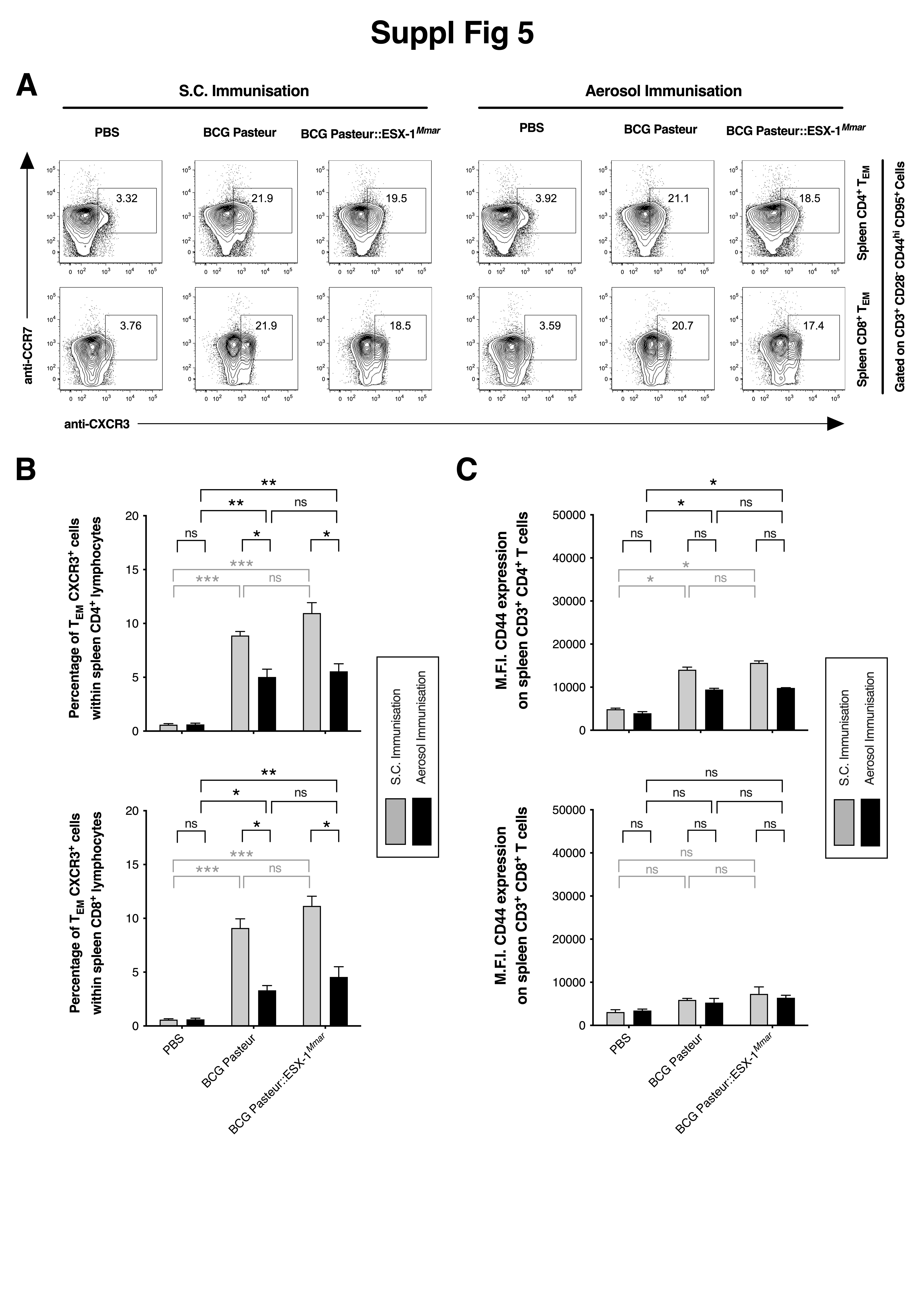

### Suppl Fig 6

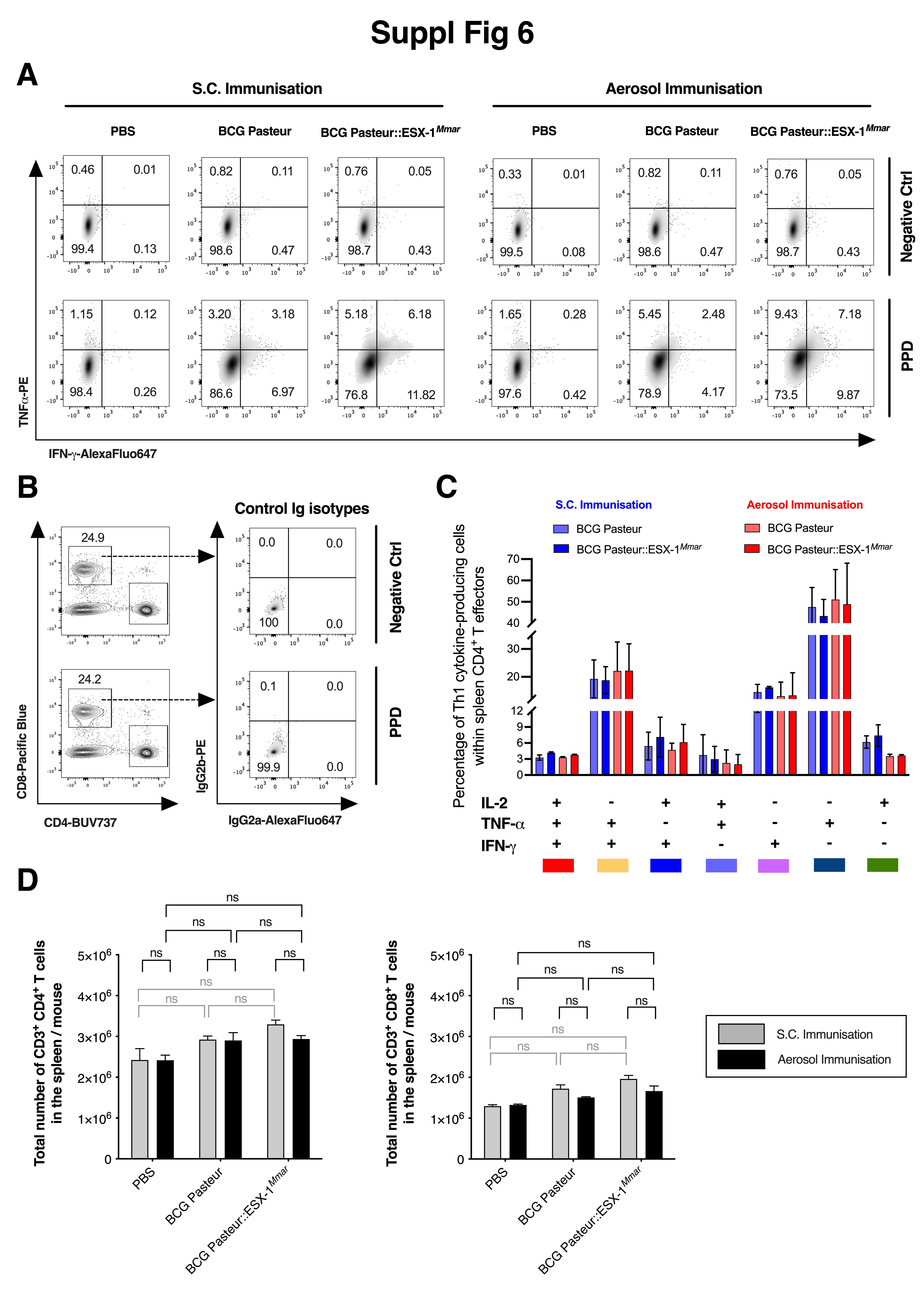

### Suppl Fig 7

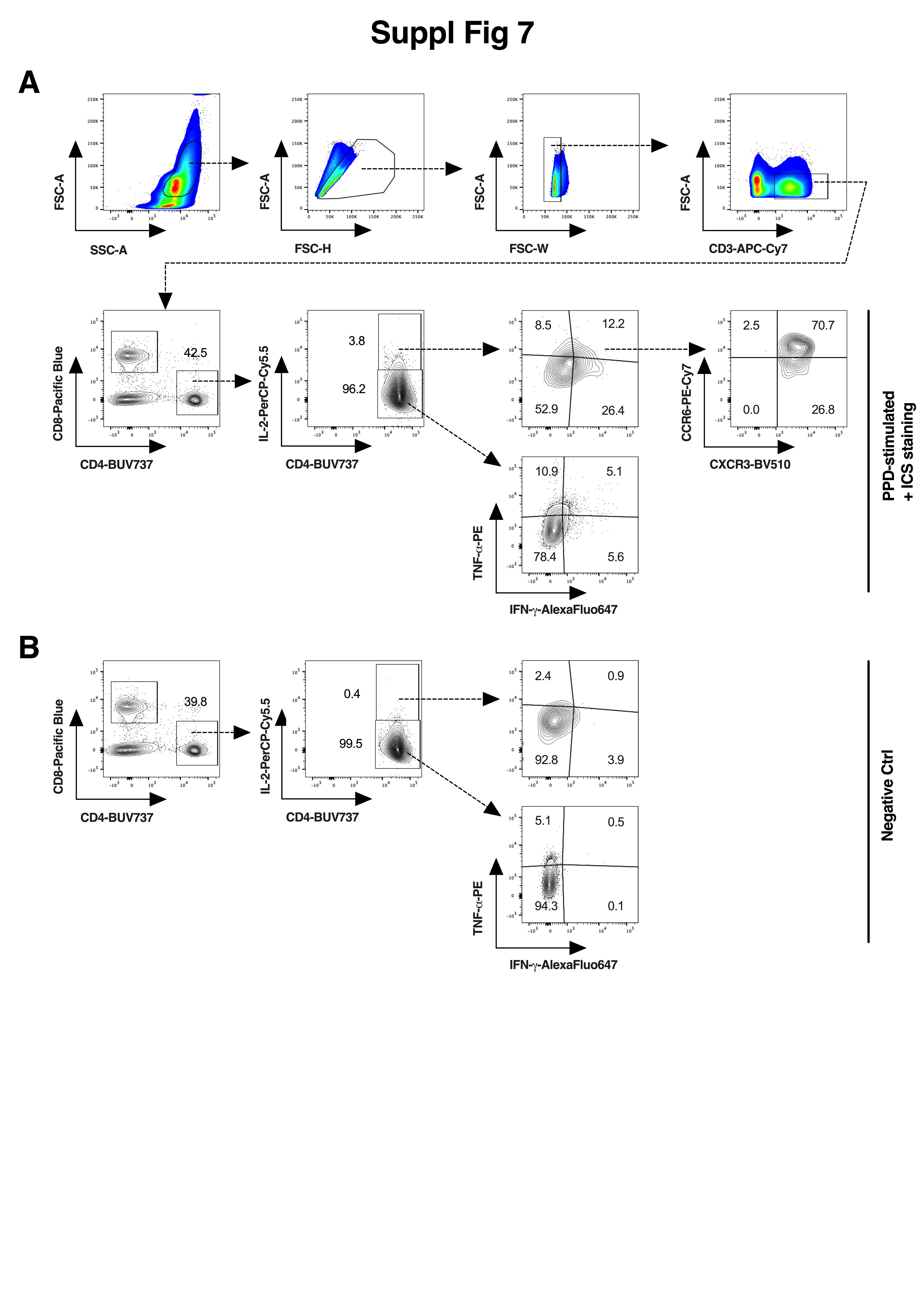
